# Reciprocal epigenetic Sox2 regulation by SMAD1-SMAD3 is critical for anoikis resistance and metastasis in cancer

**DOI:** 10.1101/2022.01.11.475900

**Authors:** Zainab Shonibare, Mehri Monavarian, Kathleen O’ Connell, Diego Altomare, Abigail Shelton, Shubham Mehta, Renata Jaskula-Sztul, Rebecca Phaeton, Mark D. Starr, Regina Whitaker, Andrew Berchuck, Andrew B Nixon, Rebecca Arend, Nam Y Lee, C. Ryan Miller, Nadine Hempel, Karthikeyan Mythreye

## Abstract

Growth factors in the tumor environment are key regulators of cell survival and metastasis. Here we reveal, dichotomy between TGF-β superfamily growth factors BMP and TGF-β/activin, and their downstream SMAD effectors. Gene expression profiling uncovered Sox2 as a key signaling node regulated in an opposing manner by anoikis-promoting BMP2, 4 and 9, and anoikis-suppressing TGF-β and activin A. We find that Sox2 repression by BMPs robustly inhibits intraperitoneal tumor burden and increases survival in multiple ovarian cancer models. Repression of Sox2 is driven by SMAD1 dependent histone H3K27me3 recruitment and DNA methylation at *SOX2’s* promoter. Conversely, TGF-β and activin A promote Sox2 expression, and anoikis resistance by SMAD3 mediated histone H3K4me3 recruitment. We find that balancing Sox2 levels is critical for anoikis, as transcriptomics reveals regulation of key cell death pathways. Moreover, BMP-driven SMAD1 signaling can override TGF-β and activin’s effect on Sox2, which has clinical significance due to the high levels of TGF-β we find in ovarian cancer patients. Together, our findings identify Sox2 as a contextual and contrastingly regulated key node, downstream of TGF-β superfamily members controlling anoikis and metastasis in ovarian cancers.

**Highlights:**

- Sox2 is a key node for anoikis resistance in cancer
- Sox2 is differentially regulated by TGF-β/activin and BMPs in broad cancers
- BMP9 is a robust metastasis suppressor by lowering Sox2
- Sox2 regulation is contextual, epigenetic and at the transcriptional level

## Introduction

Ascites are the accumulation of fluid in the abdomen associated with diseases of the peritoneal cavity, such as cirrhosis and abdominal tumors, with thirty percent of cases being related to ovarian cancer (OVCA). More than 90% of stage III and IV OVCA patients present with malignant ascites, which harbor clusters of cancer cells in suspension that directly contribute to metastasis[1] and are anoikis resistant. The ascites fluid that supports cell survival is enriched in growth factors that contribute to cancer recurrence and therapy resistance[2, 3]. Thus, defining specific growth factors that promote survival in the ascites and conversely defining strategies that disrupt survival and anoikis resistance will improve our ability to control recurrence and mortality of advanced OVCA patients.

The transforming growth factor-β (TGF-β) family of cytokines, consisting of TGF-βs, bone morphogenetic proteins (BMPs; also known as growth and differentiation factor [GDFs]), activins, inhibins (INHs), glial-derived neurotrophic factors (GDNFs), and Nodal[4] have crucial roles in cancer and development^8^ . Their cellular responses are initiated upon ligand binding to the type I, type II and type III cell surface TGF-β receptors. TGF-β receptors type I and II are serine threonine kinases, which form homomeric and heteromeric complexes upon ligand binding and activation [5] to phosphorylate intracellular receptor-regulated SMADs (R-SMADs). R-SMADs complex with the common SMAD4 and accumulate in the nucleus to regulate gene expression[6].

While members of the TGF-β superfamily share some similarities in the order that signaling events are orchestrated after ligand binding, they differ in their specific affinity for the receptors[5, 7] and receptor complexes that form. The BMP ligands (BMP2, BMP4, BMP9/*GDF2*, BMP10) bind to type I receptors: ALK1 (ACVRL1), ALK2 (ACVR1), ALK3 (BMPR1A), or ALK6 (BMPR1B), which recruit the type II receptor (BMPR2) leading to phosphorylation and translocation of the SMAD1/5/8-SMAD4 complex into the nucleus. BMP2 and BMP4 share a high degree of sequence identity and effectively bind ALK2, 3 and 6 receptors[8]. BMP9 and BMP10 both have high affinity for the ALK1 receptor[9]. However, only BMP9 can also bind ALK2 and ALK3/6 receptors[10]. TGF-βs and activin, on the other hand, first bind to the type II receptor (TβRII/ACTR2), which forms complexes with the TGF-β-type I receptors ALK4 (ACVR1B), ALK5 (TGFBR1), and ALK7 (ACVR1C) to mediate downstream signaling via SMAD2/3^18^. BMPs can also induce SMAD2/3 signaling via ALK3/6 (BMP-binding type I) and ALK5/7 (TGF-β-binding type I) receptors[11]. Similarly, TGF-β1 can also lead to phosphorylation of SMAD1/5 via ALK2/3 and ALK5 receptors [12]. SMAD1/5 activation by activin is seldom seen but has been reported[13].

In addition to the cross utilization of receptors, TGF-β, activin and BMP can both cooperate and antagonize each other during development and disease [14], [15], [16]. In cancer, this antagonism has been noted in glioblastomas[17] and breast cancer[18]. Indeed TGF-β and BMPs appear to have highly contextual roles in cancers, with both tumor suppressive and tumor promoting effects reported[19, 20]. Despite the known significance of the TGF-β superfamily members on signaling and cellular outcomes, no study has thus far delineated the function and relationship between the TGF-β members, including BMP2, 4, 9 and 10, TGF-β1 and activin during metastasis. Here, we set out to comprehensively delineate their functional relationship in a singular context of OVCA anoikis impacting metastasis to elucidate their pathological and signaling relationship and in doing so, identified Sox2 as a central regulated node downstream of BMP9, BMP2, BMP4, TGF-β1 and activin.

Sex-determining region Y-box 2 (Sox2), a single-exon transcription factor characterized by its high-affinity HMG-box DNA binding domain is essential during development[21] with an established reciprocal regulatory relationship with a subset of BMPs (BMP4) in development[22, 23]. The relationship between BMPs and Sox2 in cancer is less. TGF-β1, on the other hand has been shown to induce Sox2 expression in melanoma[24] and glioma[17]. Overexpression of Sox2 is a prognostic indicator in multiple cancers[25], [26]. However, how Sox2 is precisely regulated by most of the TGF-β members, and the unified contextual significance and relationship of this regulation remains poorly delineated.

We demonstrate Sox2 as a central repressed target downstream of BMP9, BMP2 and BMP4 leading to suppression of anoikis and metastasis. Sox2 repression occurs through epigenetic mechanisms mediated directly by SMAD1/5 leading to increased anoikis. Conversely, we demonstrate that TGF-β and activin, which are significantly elevated in patient ascites fluid, increase Sox2 expression in an epigenetic and SMAD3-dependent manner leading to anoikis resistance. Notably, the presence of BMPs and SMAD1 signaling can override the effects of TGF-β and activin on Sox2 regulation. Our findings implicate the use of a subset of BMPs as a therapeutic strategy and demonstrate the central role of context specific Sox2 regulation in controlling anoikis sensitivity and metastasis in ovarian cancer.

## Results

### BMPs promote tumor cell anoikis and suppress ovarian cancer cell survival and metastatic growth in the peritoneal cavity

We previously demonstrated promoter methylation of the gene for BMP9 (*GDF2*), with BMP9 increasing anoikis in a subset of cell lines in vitro[27]. To determine whether other BMP members besides BMP9 promote anoikis sensitivity, we examined the effect of BMP2 alongside BMP9 in a broad panel of OVCA cell lines. Cell lines representing a broad spectrum of OVCA including PA1 (teratocarcinoma of the ovary), OVCAR420 (ovarian serous adenocarcinoma), OVCAR3 (ovarian carcinoma high grade serous), SKOV3 (ovarian carcinoma non-serous), and OVCAR433 (ovarian serous adenocarcinoma) were grown under anchorage-independent conditions. Treatment with BMP2 and BMP9 significantly decreased the live-dead cell ratio in spheroids (1.8-4.25 times decrease, Fig. 1a). Spheroids treated with either BMP2 or BMP9 also exhibited reduced 3D invasion capabilities (55-67% reduction, Supplementary Fig. 1a). We previously reported no effect of BMP9 on in vitro cell growth and consistent with our prior findings[28], both BMP2 and BMP9 did not alter cell growth over a period of 3 days (Supplementary Fig. 1b). These data indicate that both BMP2 and BMP9 promote anoikis sensitivity and diminished spheroid invasion in a spectrum of OVCA cell lines.

**Fig. 1:**
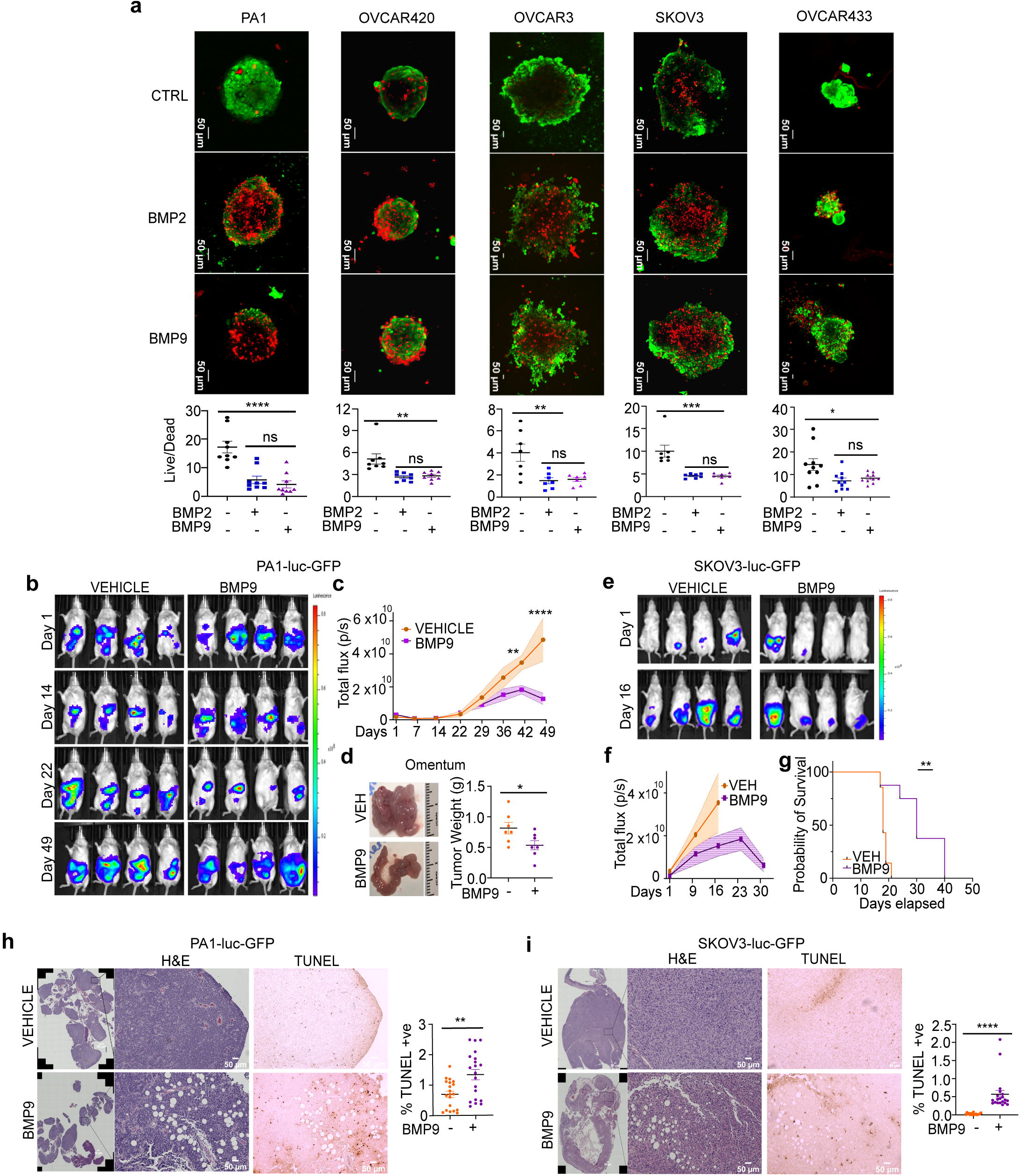
BMP’s induce anchorage-independent cell death and suppress OVCA growth and metastasis *in vivo*. **a** Representative image of OVCA cell lines cultured under anchorage independence for 48 hrs, and subsequently treated with either vehicle (VEH) control or with 10nM BMP2 or BMP9 for 24 hrs. Live/dead cell ratios were assessed by staining with Calcein-AM (green=live cells) and Ethidium homodimer dye (red=dead cells) and images taken by confocal microscopy. (Scale bar = 50µm; n=7-10). **b** Representative tumor luminescence images of NOD-SCID mice injected with PA1-luc-GFP cells either with vehicle or rhBMP9 (5mg/kg) administered daily intraperitoneally. Indicated days post-tumor cell injection from 4 mice are shown. **c** whole animal luminescence quantified over time (n=8 for rhBMP9, n=7 for vehicle). **d** Representative image of omental tumor burden (left) and quantification of omentum tumor weight (right) from mice which received either vehicle or rhBMP9 injected with PA1-luc-GFP tumor cells (n=8 for rhBMP9, n=7 for vehicle). **e** Representative tumor luminescence images of NOD-SCID mice injected with SKOV3-luc-GFP cells either with vehicle or rhBMP9 (5mg/kg) administered daily intraperitoneally. Day 1 and 16 post-tumor cell injection from 4 mice are shown. **f** whole animal luminescence quantified over time (n=8 for rhBMP9, n=7 for vehicle). **g** KM plot demonstrates increased survival of SKOV3-Luc-GFP-injected mice receiving rhBMP9 compared to vehicle. **h-i** Representative H&E and TUNEL staining (left) of (**h**) PA1-luc-GFP and (**i**) SKOV3-luc-GFP tumors demonstrate decreased tumor burden in the omentum and increased apoptosis in rhBMP treated groups compared to vehicle. TUNEL stain quantification is shown for 2 mice per group per cell line from 20 random fields/section (right; \**p* < 0.05, \*\**p* < 0.01, \*\*\**p* < 0.001). All data are presented as mean ± SEM; \**p* < 0.05, \*\**p* < 0.01, \*\*\**p* < 0.001; (**a**) two-way ANOVA followed by Dunnett’s multiple comparison test; and (**b-i**) unpaired Student’s t test.

Given the effect of BMPs (BMP2 and BMP9) on anoikis sensitivity and the established significance of anoikis in cell survival and intraperitoneal metastasis in OVCA^2^, we evaluated the effect of administering recombinant human (rh) BMP9 on peritoneal tumor growth and metastasis *in vivo*. Epithelial cancer cells express low levels of *GDF2/*BMP9[27], and delivery of BMP9 was used to mimic a potential therapeutic regimen. Overall toxicity of administering BMP9 intraperitoneal injections was examined by body weight assessment and kidney and liver function tests with no notable toxicity noted with daily BMP9 administration for up to 3 weeks (Supplementary Fig. 1c, d).

The effect of BMP9 on *in vivo* tumor growth was tested by injecting PA1 cells with either vehicle or BMP9 into the peritoneal cavity of NOD-SCID mice, followed by daily BMP9 or vehicle administration. Using bioluminescence imaging a significant reduction in overall tumor burden over time was observed in the peritoneal cavity in mice receiving BMP9 compared to mice receiving vehicle alone (Fig. 1b, c). Mice euthanized upon morbidity at the end of a 7-week study period for PA1 cells confirmed extensive tumor burden in the omentum (Fig. 1d) in vehicle treated group. In contrast, rhBMP9-treated mice had significantly lower tumor burden in the omentum (Fig. 1d). Similar results were observed with a second tumor cell line SKOV3 (Fig. 1e - f). BMP9 treatment led to significantly lower intraperitoneal tumor growth as demonstrated by broad luminescence in the abdomen by day 16 in vehicle group (Fig. 1e - f). BMP9 administration significantly prolonged survival in mice compared to the vehicle group (Fig. 1g). While all vehicle mice succumbed to disease between days 17 and 21 (Fig. 1g), BMP9 treated mice survived significantly longer for between 30-40 days (Fig. 1g). Additionally, while all vehicle treated mice had some ascites, none of the rhBMP9 treated mice had any detectable ascites.

Histological comparison of tumors from both cell lines (PA1 and SKOV3) revealed tumor cells in large nodules in the omentum of vehicle treated group. In contrast, tumors from BMP9-treated mice consisted of smaller nodules with visible adipocytes (Fig. 1h, i). Apoptosis analysis by TUNEL staining also revealed an increase in TUNEL positive cells in tumors from BMP9-treated mice as compared to vehicle treated mice in both PA1-luc-GFP and SKOV3-luc-GFP groups (Fig. 1h, i. 2x increase in PA1 and 14x in SKOV3). The increases in apoptosis (TUNEL staining) and necrotic lesions (Supplementary Fig. 1e) found in tumors from BMP9-treated mice was noted widely. ELISA analysis confirmed elevated BMP9 levels in the plasma from mice (PA1 cells injected) verifying their presence in circulation (Supplementary Fig. 1f). These data demonstrate that BMP2 and BMP9 induce anoikis in OVCA cells. Moreover, the addition of (rh) BMP9 to IP-injected tumor cells *in vivo*, which mimics the shedding of tumor cells into the peritoneal cavity during metastasis, suppresses intra peritoneal tumor spread and growth and prolongs overall survival of mice in two OVCA IP-xenograft models.

### Sox2 is a repressed transcriptional target of BMP2, 4, and 9 but not BMP10 in cancer

To identify critical factors driving anoikis and tumor burden in response to BMP exposure, we compared the transcription profile of 48,226 genes in PA1 cells cultured under anoikis-inducing growth conditions treated with either BMP9 or vehicle control (Supplementary Fig. 2a). Gene expression analysis revealed 543 differentially expressed genes using a 2-fold change criterion (p<0.05, GEO deposition #GSE185924), which were further divided into upregulated (n=333) and downregulated genes (n=210) in response to BMP9 (Fig. 2a; Source Data 1). REACTOME analysis identified 18 pathways significantly altered in response to BMP9, including BMP signaling pathways, TGF-β signaling, and transcriptional regulation of pluripotency associated stemness genes (Supplementary Fig. 2b; Source Data 1). Notably, examination of the 30 top altered genes (15-up and 15-down) revealed Sox2, IGFBP5 and HTR1D as the most repressed genes in BMP9 treated cells (12.37–20-fold change in gene expression; Fig. 2b). Amongst the top 30 altered genes 28 were linked to Sox2 through PubMed literature searches^32,33,42–44,34–41^ and the GENECARD human gene database^45,35^.

**Fig. 2:**
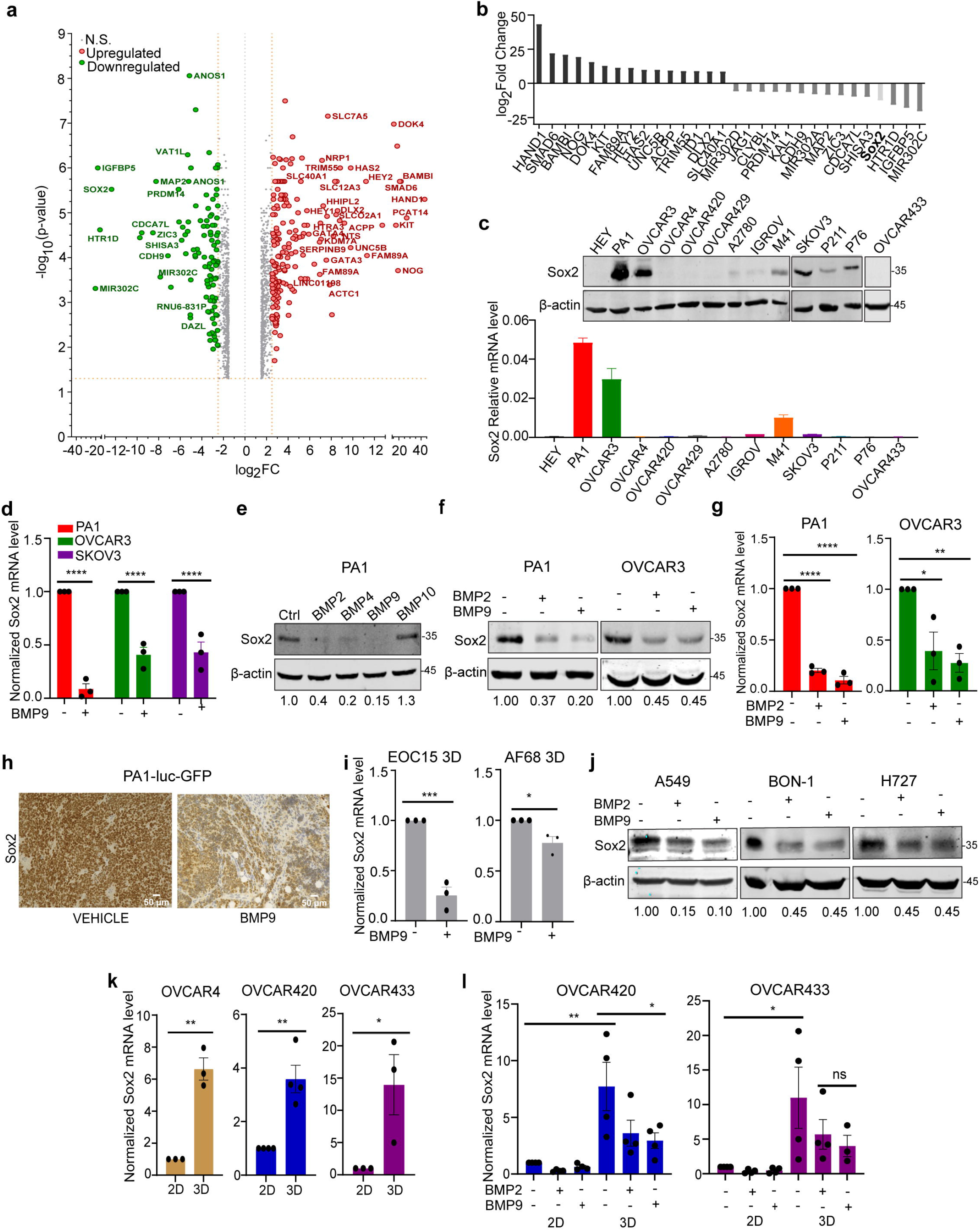
Sox2 is downregulated by BMP2, 4 and 9 in cancer cell lines and in xenograft tumors. **a** Volcano plot of changes in global gene expression in PA1 cells under anchorage-independence either treated with vehicle or rhBMP9 (cutoff 1.5 log_2_FC. Genes above cutoff of 5 are labeled). **b** List of 15 top up-regulated and down-regulated genes in response to BMP9 from (**a**). **c** Western blot (top) and qRT-PCR (bottom) screening of Sox2 expression in a panel of OVCA cells. **d** qRT-PCR analysis of Sox2 mRNA levels in response to 24 hrs BMP9 (10nM) treatment expressed relative to control untreated cells (ANOVA followed by Sidak’s multiple comparison test). **e** Western blot following treatment with BMP2,4,9 and 10 (10nM) or control for 24 hrs in PA1 cells to assess Sox2 protein expression (n=3). Quantitation of Sox2 relative to actin presented below. **f** Western blotting confirming that BMP2 and BMP9 treatment decreases Sox2 protein expression in PA1 and in OVCAR3 cells as well (n=2). Quantitation of Sox2 relative to actin presented below. **g** qRT-PCR analysis of relative Sox2 transcript levels after BMP2 and BMP9 treatment normalized to untreated control. **h** Representative immunohistochemistry (IHC) of Sox2 protein in PA1-Luc-GFP tumors from mice receiving either vehicle or rhBMP9 from Fig 1b (n= 2 mice/condition). **i** qRT-PCR analysis of relative Sox2 transcript in patient ascites-derived tumor cells maintained in ULA conditions either untreated or treated with BMP9 and normalized to untreated. **j** Western blot for Sox2 following treatment with BMP2 or BMP9 or control in cell lines of different cancer origin (n=2, A459 n=3). Quantitation of Sox2 relative to actin presented below. **k** qRT-PCR analysis of relative Sox2 expression increases in anchorage-independent (3D) conditions compared to attached (2D) culture conditions. **l** qRT-PCR analysis of relative Sox2 expression either untreated or treated with BMP2 or BMP9 under anchorage-independent (3D) conditions or attached (2D) conditions. Data are normalized to untreated attached (2D) conditions in indicated cells for **k-l**. Data are presented as mean ± SEM, \**p* < 0.05, \*\**p* < 0.01, \*\*\**p* < 0.001 (ANOVA followed by Dunnett’s multiple comparisons test).

In OVCA cell lines, Sox2 expression at baseline is variable, with PA1 cells expressing the highest relative mRNA and protein level of Sox2, followed by OVCAR3 and SKOV3 (Fig. 2c). Using this panel as a guide, we validated Sox2 downregulation by BMP9 via qRT-PCR in anchorage independent conditions (Supplementary Fig. 2c). BMP9 did not have any significant effect on two other developmental transcription factors Nanog or Oct4 (Supplementary Fig. 2d). Reduction in Sox2 expression by BMP9 was significant in several cell lines even under attached growth conditions as well including OVCAR3 and SKOV3, which express detectable RNA and protein levels of Sox2 (Fig. 2c, d). Furthermore, downregulation of Sox2 was mediated by two additional BMP ligands, BMP2 and BMP4 (Fig. 2e), with decreases at the protein (Fig. 2f) and RNA levels (Fig. 2g) in both PA1 and OVCAR3 cells. We also find that BMP4 promotes anoikis sensitivity as treatment with BMP4 significantly decreased the live-dead cell ratio in PA1 spheroids (1.9x Supplementary Fig. 2e). However, BMP10 which exhibits the highest sequence homology to BMP9^47^ did not alter Sox2 (Fig. 2e), or anoikis sensitivity (Supplementary Fig. 2f). BMP9-medaited Sox2 repression also occurred in xenograft tumors, as evaluated by Sox2 IHC in tumors obtained from vehicle or BMP9 treated mice (Fig. 1b). IHC and qRT-PCR analysis revealed an overall reduction in Sox2 levels in tumors from BMP9-treated mice compared to vehicle control-treated mice (Fig. 2h, Supplementary Fig. 2g). Importantly, we found that patient ascites-derived tumor cells, EOC15 and AF68 express Sox2 under anchorage independent conditions, which was downregulated by BMP9 treatment (Fig 2i). In addition to OVCA, several other cancer types are known to express Sox2, including lung cancer^48^, pancreatic neuroendocrine tumors, and bronchial carcinoid tumor^49,50^ . We found that BMP2 and BMP9 treatment leads to downregulation of Sox2 expression in A549 (Lung cancer), BON-1 (P-NET), and H727 (Bronchial carcinoid tumor) cells as well (Fig. 2j).

Since we observed that BMP’s induce anoikis sensitivity in both high Sox2-expressing cell lines (PA1, OVCAR3, SKOV3) and low Sox2-expressing cell lines (Fig. 1a, 2c), we tested if Sox2 levels were altered under anchorage independence, to potentially explain why BMP induces anoikis sensitivity in endogenously low Sox2 expressing cells. Indeed, we found that OVCAR4, OVCAR420 and OVCAR433 cells (low Sox2 expression) significantly upregulate their Sox2 expression under anchorage independence (Fig. 2k). These increases in Sox2 under anchorage independence were effectively suppressed by both BMP2 and BMP9 (Fig. 2l). We noted that changes in Sox2 expression under anchorage independence were not restricted to low Sox2 cell lines but also apparent in cell lines with higher baseline levels of Sox2, including OVCAR3 and PA1, which was again suppressed by BMPs (Supplementary Fig. 2c, 3a, b). Altogether, these results indicate that BMP 2, 4 and 9 can downregulate Sox2 in multiple cancer types, either when endogenous levels are high, or when Sox2 expression is induced in response to anchorage independence or during *in vivo* tumor progression.

### Rapid transcriptional regulation of Sox2 is sufficient for anoikis sensitivity

Kinetics and dose response studies reveal that repression of Sox2 by both BMP2 and BMP9 occurs in a dose dependent manner, with the effects of BMP9 being pronounced at lower equimolar doses compared to BMP2 in both PA1 and OVCAR3 cells (Fig. 3a). Sox2 protein repression began as early as 6 hours post BMP2 and BMP9 treatment and by 2 hours at the mRNA level in both PA1 (Fig. 3b) and OVCAR3 cells (Fig. 3c). We find that both BMP2 and BMP9 significantly reduce luciferase activity of a 1kb of *SOX2* promoter reporter [29] as well (Fig. 3d). Expressing Sox2 from a heterologous promoter (CMV-Sox2) prevented the BMP2 and BMP9-mediated decreases of endogenous Sox2 in SKOV3 cells (Fig, 3e). Similar overexpression of Sox2 from a second heterologous promoter (EF1a) in PA1 cells that express high levels of endogenous Sox2, was able to suppress the decrease of Sox2 by BMP2 and BMP9 (Fig, 3f) accounting for both the endogenous and heterologous EF1a driven Sox2. Overexpression of Sox2 from a heterologous promoter resulted in decreased anoikis sensitivity and more compact spheroids compared to control cells (Fig. 3g, CMV-Sox2). BMPs however had no significant effect on anoikis in cells overexpressing Sox2 from the CMV promoter (CMV-Sox2; Fig. 3h) as compared to in control cells (SKOV3-CMV-control; Fig. 3h). These data demonstrate that Sox2 confers anoikis resistance and is a major target of BMP mediated transcriptional repression.

**Fig. 3:**
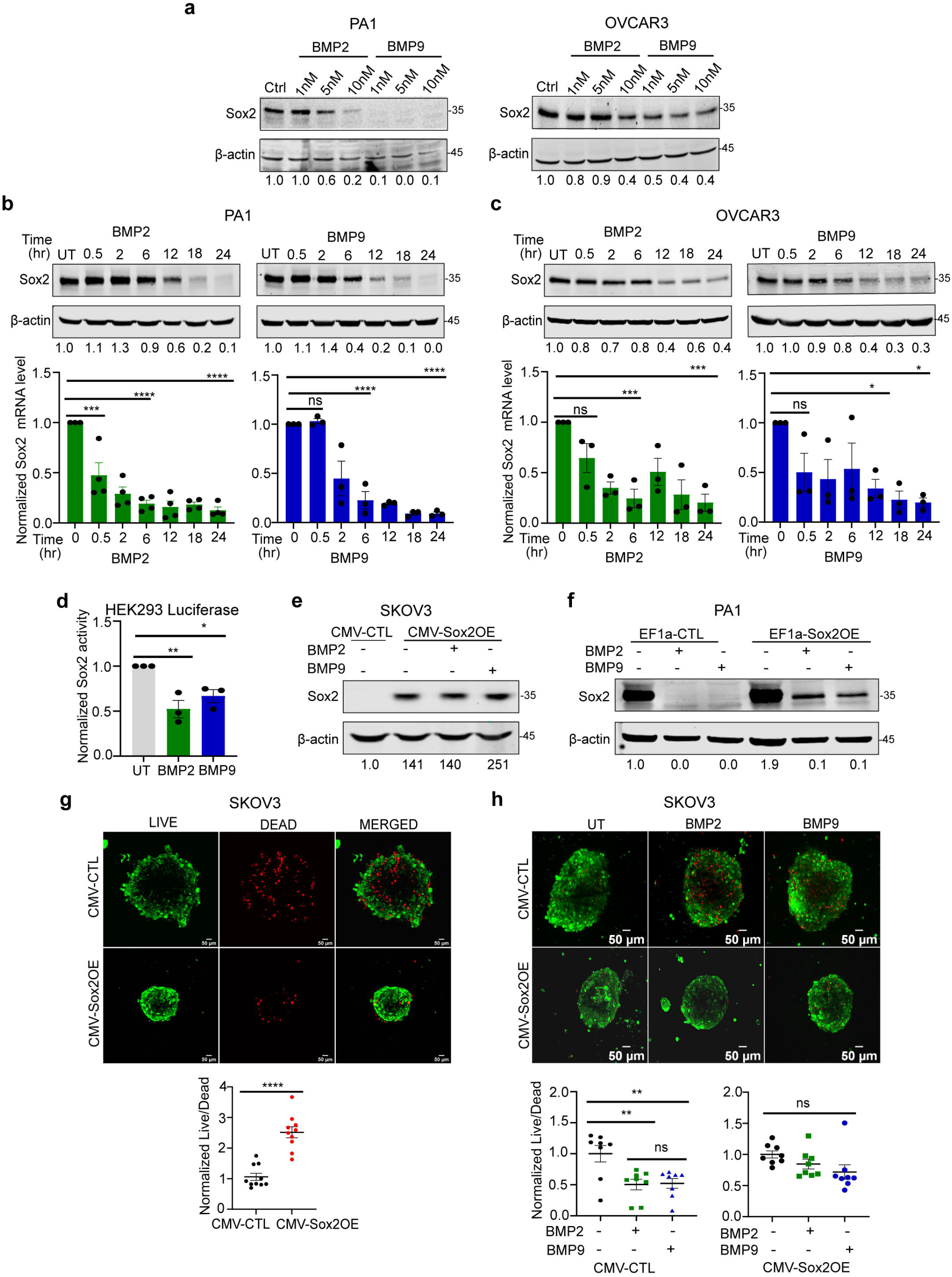
Transcriptional downregulation of Sox2 is required for anoikis sensitivity. **a** Western blot of Sox2 protein expression following BMP treatment with indicated doses for 24 hrs in PA1 and OVCAR3 cells. Quantitation of Sox2 relative to actin presented below (n=3). **b-c** Time-course analysis of Sox2 protein by western blot (top) and relative Sox2 mRNA by qRT-PCR analysis (bottom) after 10nM BMP2 and BMP9 treatment, normalized to untreated conditions (time 0 hr/UT) in (**b**) PA1 and (**c**) OVCAR3 cells. **d** pGL3-Sox2 promoter-reporter luciferase analysis in HEK293 cells following BMP2 and BMP9 treatment for 24 hrs and normalized to untreated and renilla internal control (n=3). **e-f** Western blot analysis of effect of BMP2 and BMP9 treatment for 24 hrs on Sox2 expression in CMV-CTL and CMV-Sox2 cells in (**e**) SKOV3, and EF1a-CTL and EF1a-Sox2 in (**f**) PA1 cells. **g** Representative live-dead images from SKOV3 CMV-CTL and CMV-Sox2 cells cultured under anchorage independence for 72 hrs (top) and quantified relative to CMV-TL control (bottom right. n=10; unpaired Student’s t test). **h** Representative images from SKOV3 CMV-CTL and CMV-Sox2 cells either untreated or treated with equimolar BMP2 or BMP9 for 24 hrs and live-dead ratios quantified relative to untreated control. below (n=8). Data are presented as mean ± SEM, \**p* < 0.05, \*\**p* < 0.01, \*\*\**p* < 0.001 (ANOVA followed by Dunnett’s multiple comparisons test).

### Patient ascites are low in BMP9, but high in TGF-β ligands which upregulate Sox2 and suppress anoikis

To evaluate the levels of BMPs and other TGF-β members, particularly (TGF-β1/2), in OVCA patient ascites, an environment bearing tumor cells under anchorage independence[30], we used ligand specific ELISAs. BMP9 could not be detected in patient ascites irrespective of disease stage or progression (Fig. 4a). In contrast, significantly higher levels of TGF-β1 (3800-52,348 pg/mL) and TGF-β2 (64 - 4,259 pg/mL) were detected, with TGF-β1 being an order magnitude higher than TGF-β2 (Fig. 4a). Based on these observations and the sometimes-overlapping roles of BMPs and TGF-β ligands, we tested the effect of TGF-β1 on Sox2 expression. In contrast to BMP2 and BMP9, TGF-β1 increased Sox2 protein and mRNA expression in multiple cell lines under both attached (Fig. 4b, c, d) and anchorage-independent conditions (Supplementary Fig. 4a). Activin, another TGF-β family member, also increased Sox2 levels, like TGF-β1 (Fig. 4b). In a corollary fashion, live dead analysis of anchorage-independent spheroids treated with TGF-β1 increased the live-dead ratio in multiple OVCA cells (PA1, OVCAR420 and OVCAR3; Fig. 4e), with spheroids treated with activin demonstrating a similar trend in reduction of cell death (Supplementary Fig. 4b). Luciferase activity of the 1kb *SOX2* promoter reporter construct[29] was increased in response to TGF-β1 treatment (Fig. 4f). Since BMP9 and TGF-β1 appear to have opposing effects on Sox2 expression, we evaluated if TGF-β or activin would override the negative effects of BMP9 on Sox2 expression. Equimolar amounts of BMP9 or TGF-β either decreased or increased Sox2 respectively, while the combination treatment led to a 70% reduction in Sox2 levels (Fig. 4g). Similar lowering of Sox2 was observed with the combination treatment of activin and BMP9 (Supplementary Fig. 4c). These findings on the differential effects of BMP and TGF-β on Sox2 regulation and anoikis sensitivity (Fig. 3,4), with BMP9 being able to override the effect of TGF-βs on Sox2 expression, point to Sox2 as an important regulatory node determining anoikis sensitivity of OVCA cells in response to TGF-β ligands.

**Fig. 4:**
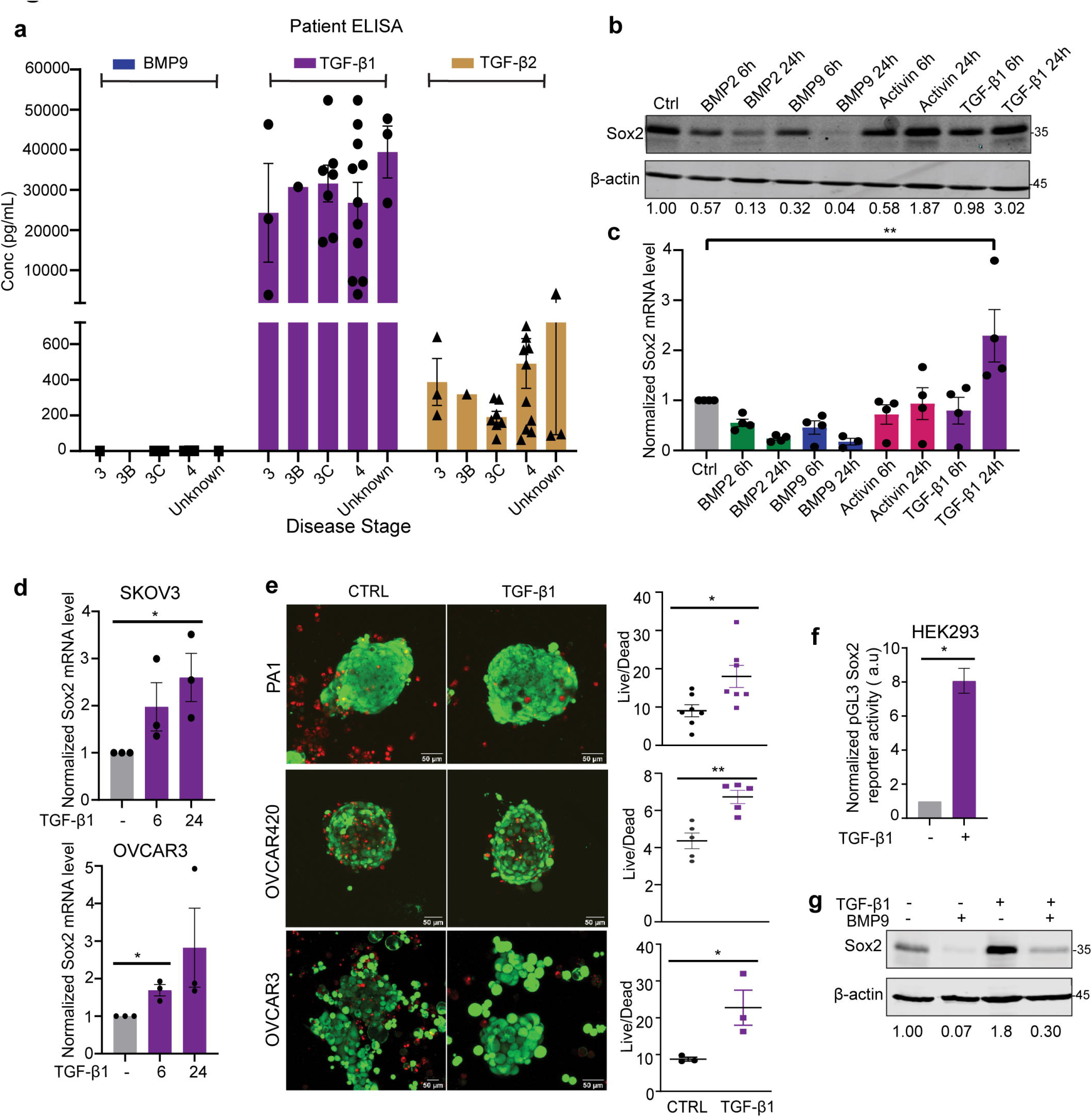
Ovarian cancer ascites are high in TGF-β ligands, which upregulate Sox2 transcription and suppress anoikis. **a** Concentration of indicated ligands in ovarian cancer patient-derived ascitic fluid (BMP9 n=10, TGF-β1 n=25, and TGFβ2 n=25) obtained using ligand specific ELISAs. **b** Western blot of Sox2 after treatment with indicated growth factors in PA1 cells. Quantitation of Sox2 relative to actin presented below (n=3). **c** qRT-PCR of Sox2 after treatment with indicated growth factors for indicated times in PA1 cells (ANOVA followed by Dunnett’s multiple comparisons test). **d** Time-course analysis of Sox2 by qRT-PCR after TGF-β1 treatment in indicated cells (ANOVA followed by Sidak’s multiple comparison test and unpaired Student’s t test). **e** Representative images from live-dead analysis upon TGF-β1 treatment in indicated cells. Quantitation of live-dead ratio in spheroid (n=3 to 7). Scale bar = 50µm. **f** pGL3-Sox2 promoter-reporter luciferase analysis upon TGF-β1 treatment for 24 hrs in HEK293 cells normalized to untreated and renilla internal control (n=3). (unpaired Student’s t test).**g** Western blot of Sox2 after combined treatment of equimolar (1nM) TGF-β1 and BMP9 for 24 hrs in PA1 cells (n=3). Data are presented as mean ± SEM, \**p* < 0.05, \*\**p* < 0.01, \*\*\**p* < 0.001.

### Sox2 levels are differentially regulated by ALK2, ALK3 and ALK5

Our findings that TGF-β ligands upregulate Sox2 while BMPs repress Sox2 expression, and that the extent of Sox2 repression differs between BMP ligand isoforms (BMP9>>BMP2, Fig. 3,4b), prompted us to further delineate the specific receptors and signaling pathways downstream of these ligands. We used a panel of small molecule inhibitors to the different Type I (ALK) receptors; Dorsomorphin (DM) to inhibit ALK2,3,6[31]; SB431542 to inhibit ALK4,5,7[32]; ML347 to inhibit ALK1,2[33] and LDN193189 to inhibit ALK2,3[34]. While BMP9 repressed Sox2 both at the protein and mRNA level in vehicle-control cells (Fig. 5a in PA1; Supplementary Fig. 5a for OVCAR3), inhibiting ALK 2,3,6 receptors resulted in 34.3% recovery in Sox2 protein levels in the presence of BMP9 (Fig. 5a DM lanes). Inhibiting ALK 4,5,7 did not significantly alter the extent of Sox2 repression by BMP (Fig. 5a SB lanes). Similarly, inhibiting ALK 2,3,6 receptors in BMP2 treated cells resulted in a 23% recovery in Sox2 repression (Fig. 5b DM lanes and Supplementary Fig. 5b; OVCAR3). Again, inhibiting ALK 4,5,7 did not alter the extent of Sox2 repression by BMP2 (Fig. 5b SB lanes). Changes to pSMAD1 were monitored in response to BMP2 and 9, and were repressed by dorsomorphin (ALK2,3,6 inhibition), with no effects observed upon co-treatment with SB431542 (ALK4,5,7 inhibition). Interestingly, inhibiting ALK 1,2 with ML347 increased baseline Sox2 protein levels even in the absence of exogenous ligand (Fig. 5c ML347 lane 4) and abrogated the ability of BMP9 to repress Sox2 at both the protein and mRNA levels in PA1 (Fig. 5c ML347+BMP9 compared to BMP9 lanes) and OVCAR3 cell line (Supplementary Fig. 5c). In comparison to BMP9, ALK1,2 inhibition only partially prevented BMP2 mediated Sox2 repression (72% recovery Fig. 5c ML347+BMP2 compared to BMP2 lanes). BMP9-induced phosphorylation of SMAD1 was completely suppressed in ML347 treated cells (Fig. 5c, ML347 lanes, Supplementary Fig. 5c), and this only partially (40%) suppressed BMP2-induced SMAD1-phosphorylation (Fig. 5c). Similarly, inhibition of ALK2,3 using LDN193189 increased Sox2 protein levels at baseline even in the absence of exogenous ligand (3x, Fig. 5c) and blocked both BMP2 and BMP9’s ability to repress Sox2 at the protein and mRNA level (Fig. 5c). Full inhibition of BMP2 and BMP9 induced SMAD1/5 phosphorylation was also observed (Fig. 5c LDN lanes, and Supplementary Fig. 5c; OVCAR3 cells). Altogether these data demonstrate a strong preference for ALK2 in mediating BMP9-dependent Sox2 repression, and a combination of ALK2 and ALK3 in driving BMP2-dependent Sox2 repression.

**Fig. 5:**
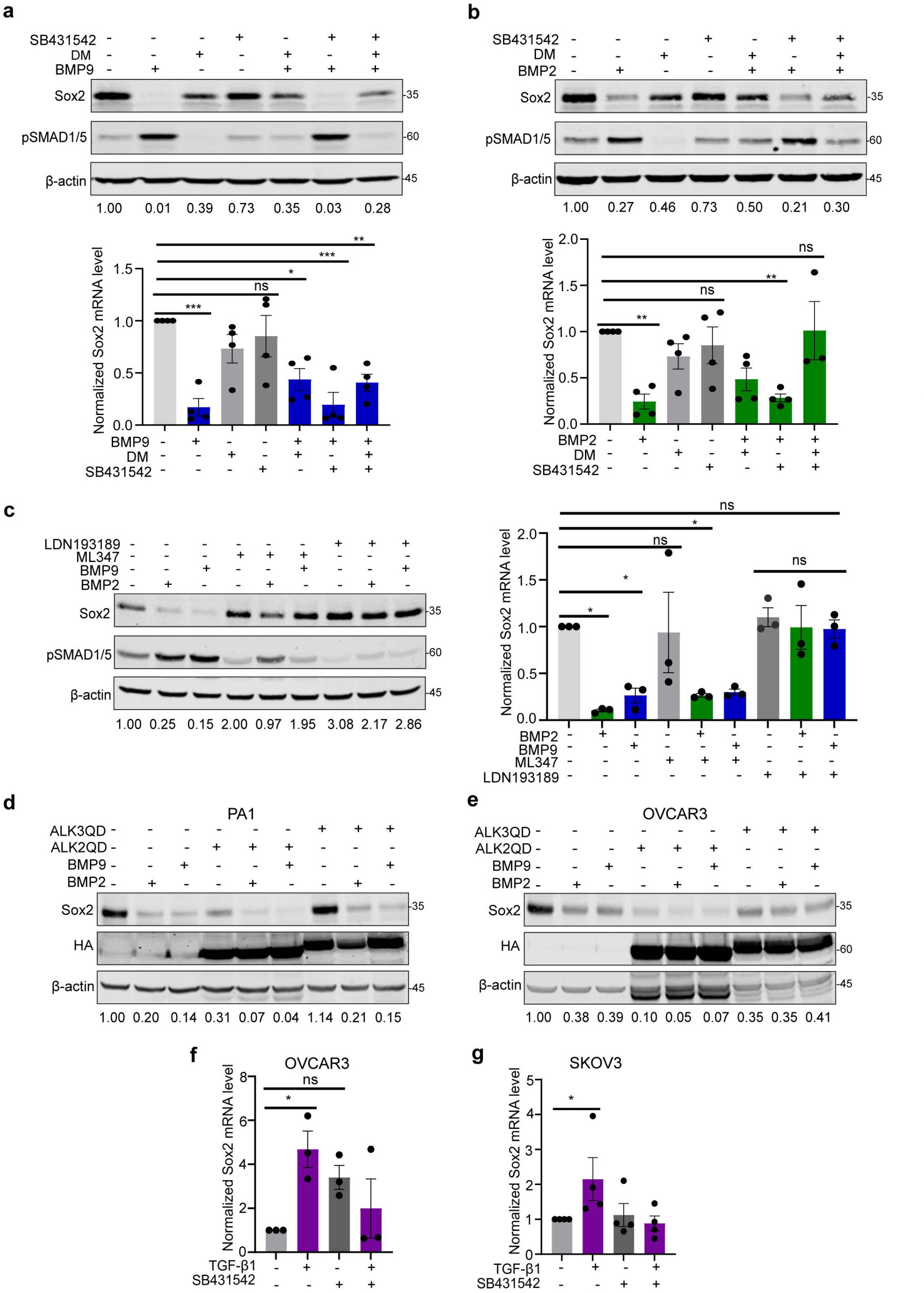
Sox2 is reciprocally regulated by ALK2/ALK3 and ALK5 receptors. **a** Western blot (top) and qRT-PCR (bottom) analysis of Sox2 expression in PA1 cells pretreated with 5μM ALK2,3,6 inhibitor Dorsomorphin (DM) and 5μM ALK4,5,7 inhibitor SB431542 for 1hr, followed by treatment with BMP9 for 24 hrs. Data are normalized to vehicle DMSO controls. WB quantitation of Sox2 relative to actin presented below. **b** Western blot (top) and qRT-PCR (bottom) analysis of Sox2 expression in PA1 cells pretreated with 5μM Dorsomorphin (DM) and 5μM SB431542 for 1hr, followed by treatment with BMP2 for 24 hrs. Data are normalized to DMSO vehicle controls. WB quantitation of Sox2 relative to actin presented below (n =4 for DM and 2 for SB).**c** Western blot (left) and qRT-PCR (right) analysis of Sox2 expression in PA1 cells pretreated with 3μM ALK1,2 inhibitor ML347 and 0.8μM ALK2,3 LDN193189 for 1hr, followed by treatment with BMP2/9 for 24 hrs. Data are normalized to vehicle controls presented. WB quantitation of Sox2 relative to actin presented below (n=2). **d-e** Western blot of Sox2 in cells expressing ALK2QD, ALK3QD or vector control after 24 hrs treatment of BMP2 and BMP9 (n=2) in (**d**) PA1 and (**e**) OVCAR3 cells. **f-g** qRT-PCR of Sox2 in indicated cells pretreated with 5μM SB431542 for 1hr, followed by treatment with TGF-β1 for 24 hrs. Data are normalized to DMSO controls presented. All sata are presented as mean ± SEM, \**p* < 0.05, \*\**p* < 0.01, \*\*\**p* < 0.001 (**a-f**) ANOVA followed by Dunnett’s multiple comparisons test and (**g**) ANOVA followed by Sidak’s multiple comparisons test.

To confirm the specific roles of ALK2 and ALK3 receptors in Sox2 repression, recombinant constitutively active kinases ALK2 or ALK3 (HA-ALK2QD and HA-ALK3QD) were expressed in both PA1 and OVCAR3 cells. Activating ALK2 kinase (ALK2QD) alone decreased Sox2 expression even in the absence of exogenous BMP ligand in both cell lines (69% reduction in PA1 and 90% in OVCAR3 Fig. 5d, e). In the presence of ligand (BMP2, BMP9), ALK2QD-mediated Sox2 repression was further enhanced (Fig. 5d, e). The effect of activating ALK3 (ALK3QD) was more modest compared to ALK2QD and was cell line dependent. ALK3QD did not reduce Sox2 in the absence of exogenous ligand in PA1 cells but was able to reduce Sox2 levels by 65% in OVCAR3 cells in the absence of exogenous ligand (Fig. 5d, e). The presence of ligand (BMP2, BMP9) only slightly enhanced Sox2 repression in both cell lines (Fig. 5d, e). These findings demonstrate a requirement for both ALK2 and ALK3, with ALK2 being critical for maximum Sox2 downregulation based on ligand independent effects and enhancement of the effects of both BMP9 and BMP2.

Since both TGF-β1 and 2, and activin predominantly utilize ALK 4,5,7 for phosphorylating SMAD2/3, we evaluated the effect of blocking their kinase activity using SB431542. We find that treatment with SB431542 suppressed TGF-β1 induced Sox2 increase (Fig. 5f, g). These studies together implicate different ALK receptors: ALK2 and ALK3 in Sox2 repression, and ALK4/5 in Sox2 activation.

### SMAD1 and SMAD3 differentially regulate Sox2 and occupy the *SOX2* promoter in response to BMP9 and TGF-β respectively

SMAD1 phosphorylation is a primary response to ALK2 and ALK3 kinases[5] that regulate Sox2 levels downstream of BMP (Fig. 5). Additionally, we previously reported a SMAD1 signaling preference for BMP9[27]. Hence, we tested a direct role for SMAD1 in Sox2 repression using pooled shRNAs to SMAD1 (shSMAD1). Reducing SMAD1 significantly decreased the ability of BMP2 and BMP9 to reduce Sox2 levels compared to control shRNA cells (shNTC) by 30-44% at the protein level (Fig. 6a). Since ALK5, the type I receptor downstream of TGF-β signaling required for increasing Sox2 expression (Fig. 5f, g) primarily phosphorylates SMAD2/3[5], we silenced SMAD3 using pooled siRNAs (Fig 6b). While TGF-β increased Sox2 levels in control (siScr) cells (Fig. 6b), TGF-β was unable to increase Sox2 in siSMAD3 transfected cells (Fig. 6b). Strikingly, siRNA to SMAD3 also lowered Sox2 levels at the baseline even in the absence of exogenous ligands (Fig. 6b), indicating direct roles for SMAD3 in Sox2 upregulation. We also observed a likely compensatory increase in pSMAD1 upon lowering SMAD3 (Fig. 6b) that correlated with lower Sox2 levels even in the presence of TGF-β1 (Fig. 6b).

**Fig. 6:**
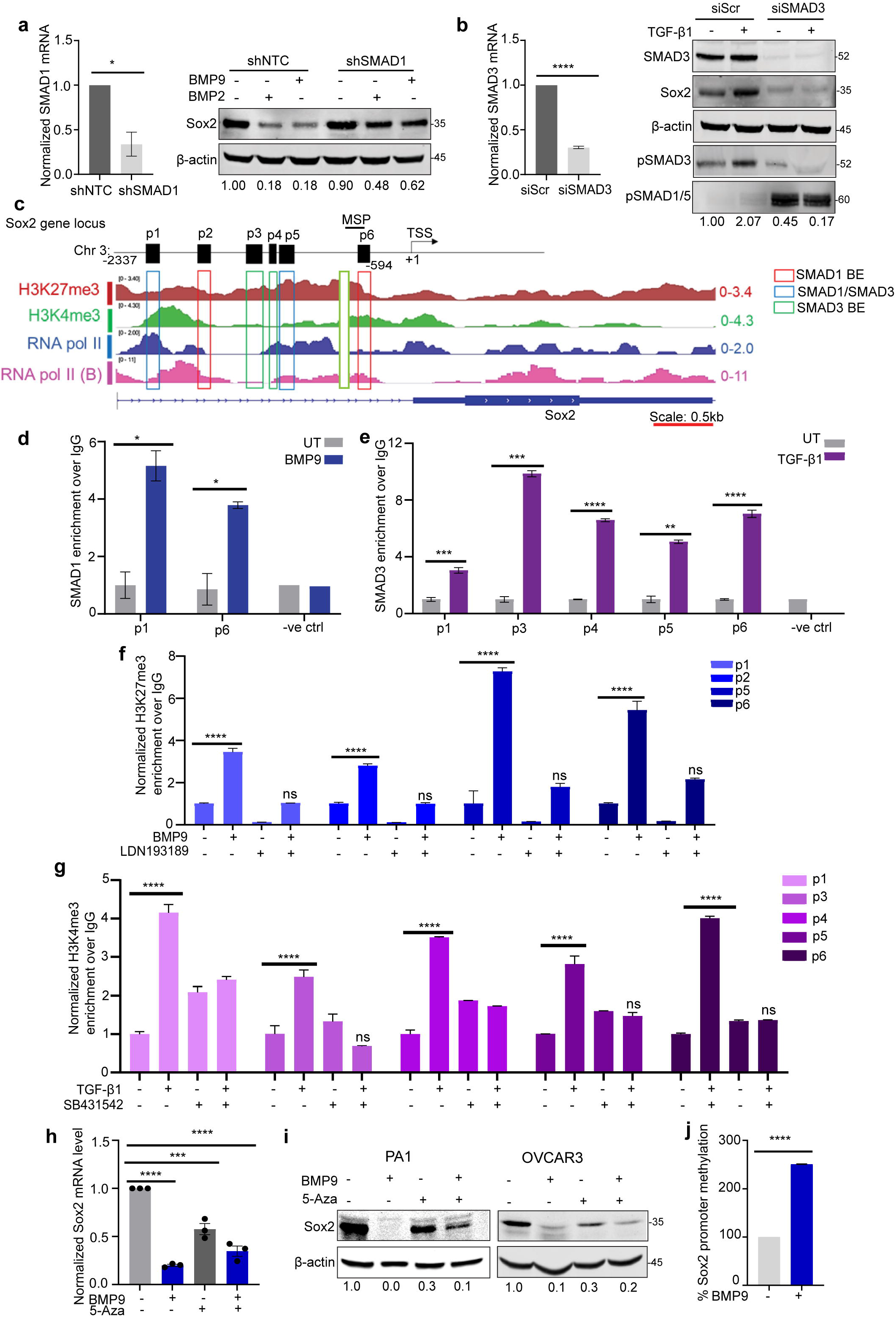
SMAD1 and SMAD3 directly regulate Sox2 expression and occupy *SOX2’s* promoter at distinct and overlapping sites. **a** Relative qRT-PCR of SMAD1 levels in shSMAD1 cells normalized to shNTC in OVCAR3 cells (left). Western blot analysis of Sox2 in OVCAR3 shSMAD1 or non-targeting control (shNTC) treated with indicated equimolar BMPs’ for 24 hrs (right). (n=3) **b** qRT-PCR analysis of SMAD3 levels in OVCAR3 cells transiently expressing siRNA to SMAD3 (siSMAD3) or scramble control (siScr). Data are normalized to siScr in OVCAR3 cells (left) (\**p* < 0.05, \*\**p* < 0.01, \*\*\**p* < 0.001 two-tailed student t test). Western blot analysis of indicated proteins in OVCAR3 siSMAD3 or scramble control (siScr) cells treated with TGF-β1 for 24 hrs with quantitation of Sox2 relative to actin presented below (right) (n=3). **c** In-silico analysis showing primer sites flanking SMAD1 and SMAD3 Binding Element (BE) in chromosomal region including Sox2’s promoter and gene as indicated. TSS= transcription start site, MSP-Methylation Specific PCR Primer. **d** Representative relative qRT-PCR of indicated regions (primer sites) after chromatin immunoprecipitation of SMAD1 to sites on Sox2 proximal chromosomal regions with or without 1hr of BMP9 treatment, expressed as the ratio over IgG controls normalized to untreated cells (n=3). **e** Representative relative qRT-PCR of indicated regions (primer sites) after chromatin immunoprecipitation of SMAD3 to sites on Sox2 proximal chromosomal regions with or without 1hr of TGF-β1 treatment, expressed as the ratio over IgG controls normalized to untreated cells (n=3). **f** Representative qRT-PCR of indicated regions (primer sites) associated with H3K27me3 enrichment to sites on Sox2 proximal chromosomal regions with and without 1hr BMP9 treatment, +/- LDN193189 as indicated in PA1 cells expressed as the ratio over IgG controls normalized to untreated cells (n=3 for BMP9, n=2 for LDN). **g** Representative qRT-PCR of indicated regions (primer sites) after chromatin immunoprecipitation with H3K4me3 to sites on Sox2 proximal chromosomal regions with and without 1 hr TGF-β1 treatment, +/- SB431542 as indicated in PA1 cells expressed as the ratio over IgG controls normalized to untreated cells (n=3 for TGF-β1, n=2 for SB). **h** qRT-PCR of Sox2 in PA1 cells pretreated with 5μM 5-Azacytidine (5- Aza) for 48 hrs, followed by treatment with BMP9 for 24 hrs, normalized to DMSO controls. Data are presented as mean ± SEM, \**p* < 0.05, \*\**p* < 0.01, \*\*\**p* < 0.001 (ANOVA followed by Dunnett’s multiple comparisons test). **i** Representative western blot of Sox2 in indicated cells pretreated with 5μM 5-Azacytidine (5-Aza) for 48 hrs, followed by treatment with BMP9 for 24 hrs. Quantitation of Sox2 normalized to actin and DMSO controls presented (n=2). **j** Representative MS-qPCR using MS qPCR primers (Fig 6c) to Sox2 proximal to Sox2’s TSS, with or without BMP9 treatment for 24 hrs normalized to untreated control (n=2). (\**p* < 0.05, \*\**p* < 0.01, \*\*\**p* < 0.001 two-tailed student t test).

In-silico analysis revealed several SMAD1 and SMAD3 binding motifs (GG(C/A)GCC and GTCT/AGAC, respectively) within 2 kb upstream of the transcriptional start site for *SOX2* (TSS; Fig. 6c)^55^. Chromatin immunoprecipitation (ChIP) was used to assess the binding of SMADs to these sites. The promoter contains four SMAD1, two SMAD3 binding motifs. However, due to several ‘CG’ clusters (CpG islands), we designed primers flanking regions immediately outside the CpG islands with additional sites within 2 Kb of the TSS (Fig 6c). Primers flanking the SMAD1 binding elements (p1, p2, p5, p6) and primers flanking SMAD3 binding elements (p1, p3, p4, and p5) were used, with p1, and p5 having both SMAD1 and SMAD3 binding elements, and p6 located closest to the TSS (Fig. 6c). BMP9 treatment led to a significant enrichment of SMAD1 binding at two sites: p1 and p6 (Fig. 6d). p2 and p5 had modest SMAD1 enrichment but were not consistent in our independent biological experimental trials and hence are not shown here. TGF-β1 treatment led to consistent SMAD3 enrichment at p1, p3, p4, and p5 (Fig. 6e). Due to the proximity of p6 to a SMAD3 binding element, we also tested and find SMAD3 enrichment at p6 in response to TGF-β1 treatment (Fig. 6e). These sites were also tested for response to activin and were similarly found to be occupied by SMAD3 (Supplementary Fig. 6a). These data together indicate enrichment of SMAD1 and SMAD3 to *SOX*2’s promoter and upstream regions in response to BMP9, TGF-β1 and activin respectively with one site occupied by both SMAD1 and SMAD3 and other uniquely occupied regions.

### Epigenetic regulation of Sox2 is mediated by SMAD-dependent methylated histone occupancy and promoter DNA methylation

Gene repression and activation by SMADs frequently requires additional proteins and chromatin modification[35]. Hence, we evaluated whether new transcription, protein synthesis or protein turnover/degradation was required for BMP9 induced Sox2 repression. We find that treating cells with cycloheximide to induce translation arrest^51^ dampened the repressive effect of BMP2/9 on Sox2 protein and mRNA (3.5-14 times and 3.6-20 times respectively, Supplementary Fig. 6b). Similarly, actinomycin D (transcription arrest) treatment blocked Sox2 mRNA downregulation by BMPs (6 times, Supplementary Fig. 6c). No effect of MG132, inhibitor of proteasomal degradation, on BMP’s ability to repress Sox2 was seen (Supplementary Fig. 6d). Taken together, these findings indicate a likely requirement of new/additional protein synthesis for repression. Notably H3K27me3 was significantly enriched on the *SOX2* promoter in response to BMP9 treatment as determined by ChIP (Fig. 6f). This enrichment was SMAD1 signaling dependent, as H3K27me3 occupancy was significantly reduced in the presence of LDN193189 (ALK/SMAD1 inhibitor; Fig. 6f). Conversely TGF-β1 treatment led to enrichment of H3K4me3 2.5-4.1x at multiple SMAD3 motifs (Fig. 6g, p1, p3, p4, p5, p6). This enrichment was SMAD3-dependent as H3K4me3 enrichment was abrogated by SB431542 (Fig. 6g). These regions are consistent with ENCODE analysis in Hela and A549 cell line (Fig. 6c) that identified highest H3K27me3 peaks at p5 and p6, the same regions we found SMAD1 enrichment in response to BMP9 (Fig. 6d). p1 exhibited a high peak of H3K4me3, the same region as SMAD3 occupancy in response to TGF-β1. Our finding suggests that SMAD1 signaling, and occupancy, leads to increases in H3K27me3 histones and conversely SMAD3 signaling, and occupancy leads to increases in H3K4me3 histones at *SOX2*’s promoter and regulatory regions.

Due to the presence of cluster of CpG islands withing10bp from the p6 primer sites, (Fig. 6c), we evaluated the effect of DNA methylation in this region in response to BMP9. We find that treatment with DNA methyltransferase (DNMT) inhibitor, 5’-azacytidine (5’-Aza) suppressed Sox2 mRNA downregulation by BMP9, resulting in a partial recovery in Sox2 expression at the mRNA and protein level in both PA1 (Fig. 6h, i) and OVCAR3 cells (Fig. 6i). Methylation specific qPCR in response to BMP9 revealed a 2.5x increase in *SOX2* promoter methylation status in response to BMP9 treatment as compared to untreated control cells (Fig. 6j). These findings indicate that DNA methylation along with SMAD1-dependent H3K27me3 recruitment are drivers of Sox2 repression in response to BMPs.

### Sox2 repression leads to genome wide changes in key transcriptional factors and cell death pathways under anchorage independence

High Sox2 is associated with a poor prognosis for OVCAs[36] and reducing Sox2 expression transiently using pooled siRNAs (siSox2) (Fig. 7a) or alternatively stably using shRNA (shSox2) (Fig. 7b) resulted in increased anoikis sensitivity under anchorage independence compared to control cells (Fig. 7a, b). In both siSox2 and shSox2 cells, spheroids appeared disaggregated and less compact compared to their respective controls (Fig. 7a, b). The critical requirement of Sox2 for anoikis resistance under anchorage independence led us to explore genes and pathways impacted by alterations to Sox2 specifically under anchorage independence. Genome wide gene expression profiles of siSox2 and control siNTC PA1 cells were compared using RNA-sequencing under anchorage independence. Our analysis revealed 59 differentially expressed genes (DEG) between siNTC and siSox2 cells (p-value ≤ 0.05, Fig. 7c; Source Data 2). Of these, 24 genes were downregulated while 35 genes were upregulated in siSox2 cells compared to control (Fig. 7c; Source Data 2). Of the total 59 genes, 21 of these including Sox2 also changed their expression levels in response to BMP9 treatment from our microarray analysis (Fig. 7d, Supplementary Fig. 7a). A closer analysis of the 20 genes for Sox2 binding motifs within one kilobase of the transcription start sites using LASAGNA-Search, revealed that 17/20 of the common DEG’s presented one or more Sox2 binding motif (Supplementary Fig. 7b). Downregulated DEGs had previously been implicated in processes relevant to cell adhesion (POSTN[37]) and metastasis (TRIM22)[38]. Gene set enrichment analysis of all the differentially expressed genes from the RNA-seq data revealed enrichment of 8 Hallmark gene sets based on an FDR value <25% and included ‘Apoptosis’, ‘TGF-β signaling’, and ‘Epithelial-Mesenchymal transition’ in siSox2 cells (Fig. 7e) and interestingly, ‘interferon alpha response’ and ‘interferon gamma response’ in the control siNTC cells (Fig. 7f). Upregulation of several genes from the Apoptosis pathway, including BMF, BCL2 L11 and BID (Fig. 7e) were confirmed to be upregulated in siSox2 cells compared to control cells under anchorage independence (Supplementary Fig. 7c). Moreover, we also analyzed genes from the ‘TGF-β signaling’ pathway and identified ACVR1 (ALK2) as one of the upregulated genes upon silencing of Sox2 (Supplementary Fig. 7c). Additional validated genes from the RNA seq analysis included TRIM22, CD47 and CD74 genes from the interferon alpha and gamma response pathways in si-control (siNTC) cells (Supplementary Fig. 7c, d). We find TRIM22 was downregulated in siSox2 cells (Supplementary Fig. 7c) while CD47 and CD74 were upregulated in siSox2 cells in PA1 and OVCAR3 (Supplementary Fig. 7d). Taken together, these findings establish a role for Sox2 silencing in promoting apoptosis under anchorage independence with potential alterations to key transcriptional and epigenetic regulators and adhesion molecules for tumor cell survival.

**Fig. 7:**
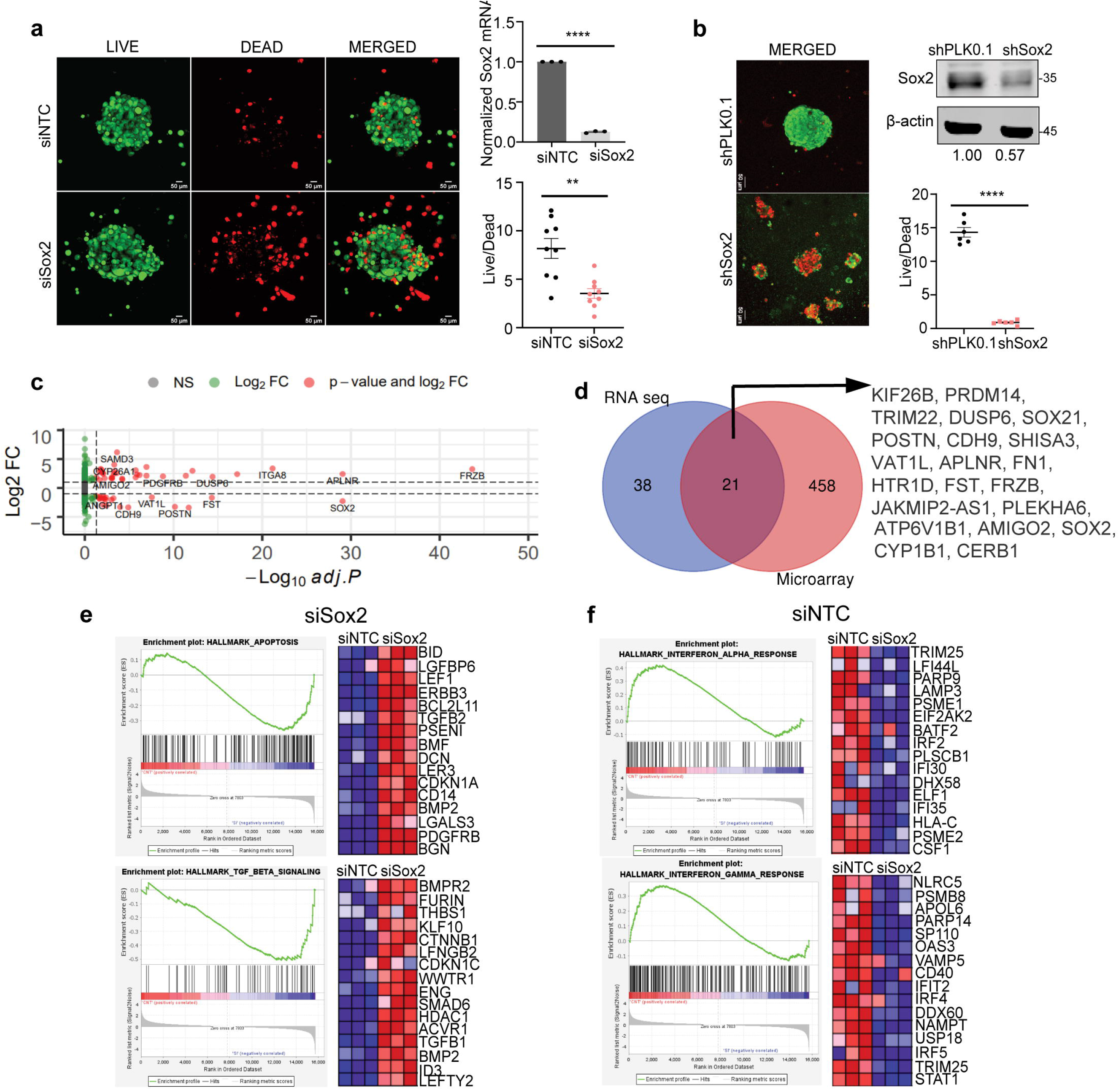
Genome-wide transcriptome changes upon reducing Sox2 and increasing anoikis, reveal apoptotic pathways and key transcriptional epigenetic regulators and adhesion molecules. **a** Representative image from siNTC or siSox2 PA1 cells under anchorage independence for 72 hrs (left). qRT-PCR of Sox2 expression in siSox2 cells normalized to siNTC cells (top right) and quantitation of live-dead ratio in spheroid cells (n=9) (bottom right). **b** Representative images from PA1 shPLKO.1 and shSox2 cells under anchorage independence for 72 hrs (left). Western blot of Sox2 expression in shPLK0.1 and shSox2 cells (top right), and quantitation of live-dead ratio in spheroid cells (n=6) (bottom right). \**p* < 0.05, \*\**p* < 0.01, \*\*\**p* < 0.001. **c** Volcano plot of significant differentially expressed genes (DEG) based on adjusted p-value of 0.05 between siNTC and siSox2 in PA1 cells under anchorage independence growth for 48hrs. **d** Venn diagram of common DEG’s between RNA seq data from (**a**) and microarray data from BMP9 treatment under anchorage independence in PA1 cells from (Fig 2a). **e-f** Gene Set Enrichment Analysis (GSEA) of pathways differentially altered in (**e**) siSox2 and (**f**) siNTC with corresponding Blue-Pink O’gram of core enrichment genes generated by GSEA (right panel).

## Discussion

Here we provide the first in depth analysis of the role TGF-β ligand family members play in OVCA transcoelomic metastasis, by delineating distinct effects of the TGF-β, BMP and activin subfamilies. Using a combination of cell lines, patient derived samples, cell line and xenograft models, we demonstrate dichotomous roles of BMPs and TGF-β/activin and their respective downstream mediators SMAD1 and SMAD3 in the regulation of anoikis and cell survival under anchorage independence and show that this is dependent on the divergent epigenetic regulation of the developmental gene *SOX2*.

We previously observed a role for BMP9 (*GDF2*) in conferring anoikis sensitivity in a subset of epithelial cell lines[27]. Here, we find that additional BMP members, including BMP2 and BMP4, promote anoikis and that spheroids generated in response to anchorage-independence are less invasive when exposed to BMPs. BMP9 was chosen to further evaluate the effects of BMPs on transcoelomic OVCA metastasis using models that recapitulate human disease spread in the peritoneal cavity. BMP9 administration at the time of IP anchorage-independent tumor cell injection to mimic the shedding of tumor cell from the primary tumor revealed that BMP9 treatment reduced transcoelomic metastasis and prolonged overall survival from disease burden. Our findings alongside prior studies on the impact of BMP9 on normalizing tumor blood vessels (Lewis lung carcinoma)[39] suggest a potential dual role of the anti-tumor properties of BMP9 on the vasculature likely via the endothelial specific TGF-β receptor ACVRL1[40], and an anoikis-stimulating effect on epithelial cells *via* the ALK2/3 receptor, as demonstrated here. Given these findings, therapeutic strategies that combine BMP9 with anti-angiogenic approaches should be further investigated to evaluate the therapeutic window and utility of BMP9 in cancer.

We find that Sox2 lies at the center of the signaling node that drives the differential effects of BMPs and TGF-β on anoikis. While both BMP2 and BMP4 also increase anoikis, BMP9 had a more potent effect at repressing Sox2 expression at lower doses. It is possible that BMP2 induced anoikis may involve additional downstream mediators, besides Sox2. BMP10 had no significant effect on epithelial cell anoikis. This can be explained by the ability of BMP9 to utilize ALK2 [27]. Consistent with this receptor specificity model, BMP10 failed to activate SMAD1/5 [27], repress Sox2, and impact anoikis in epithelial cells . Probing the receptor signaling mechanisms revealed that ALK2/ALK3 induced phosphorylation of SMAD1 is indeed critical for Sox2 repression (Fig. 5c) as a key step in BMP-mediated anoikis sensitivity. A more robust requirement of ALK2 than ALK3 was observed for Sox2 repression, which could account for higher sensitivity to BMP9 than BMP2.

Much like other BMP’s[28] it is likely that BMP9 has a contextual role in cancer. While it strongly enhanced anoikis [27], BMP9 had no negative effects on cell viability under attached conditions. Similarly, both BMP2 and BMP9 have been showed to function as potent tumor suppressors in several cancers including, but not limited to breast[18] and prostate[41] with prior conflicting studies in OVCA indicating increased tumor growth in subcutaneous models[42] that do not accurately mimic human ovarian intraperitoneal cancer spread.

We identified Sox2 repression as a key mechanism regulated by multiple BMP members (BMP2, BMP4 and BMP9, but not BMP10) and not restricted to OVCA. The implications of this regulation are likely to have tumor specific consequences, warranting further investigation in these cancers. A key finding from our work is the divergent role of TGF-β members on anoikis and SOX2 regulation. This antagonism is of particular clinical relevance as we demonstrate that patient ascites are highly enriched in TGF-β1, while BMP9 was undetectable (Fig. 4a), suggesting that OVCA cells are primarily exposed to TGF-β1 ligands that stimulate Sox2 expression and enhance survival under anchorage independence (Fig. 4b, e). While TGF-β has been previously reported to promote spheroid invasion in breast cancer[43], activin has not previously been implicated in regulating anoikis resistance in OVCA. Our findings are consistent with studies on inhibition of TGF-β1 signaling and reduced peritoneal tumor growth in OVCA[44, 45]. With accumulating preclinical and clinical evidence on the effectiveness of inhibiting TGF-β1 and activins as a therapeutic strategy in OVCA [46], these findings highlight an important new mechanism of the pro-metastatic roles of TGF-β1 and activin through the regulation of Sox2 and anoikis resistance.

Since the TGF-β1 receptor dependencies are key in the dichotomous regulation of Sox2 by TGF-β family members, alterations in receptor expression could be an important tipping point in determining the balance between SMAD1 versus SMAD3 signaling, leading to either Sox2 downregulation or upregulation, respectively. Of note, TGF-β family members, particularly TGF-β1, can also lead to phosphorylation of SMAD1/5 via ALK2/3 and ALK5 receptors[12, 47]. However, we found that SMAD3 knock-down was sufficient in abrogating TGF-β1-mediated increases in Sox2. The compensatory increase in pSMAD1 levels that correlated with lowered Sox2 suggest that shifting the balance between SMAD1 activation and SMAD3 activation regardless of the upstream ALK involved, could potentially tip the effect of exogenous TGF-β from increasing Sox2 to suppressing Sox2. The amount of ligand is likely to also play a key role in this process.

In ligand combination studies of TGF-β1/activin with BMP9, BMP9 could override TGF-β1/activin to downregulate Sox2. Since BMP9 is unlikely to be present at the same levels as TGF-β in cancer , such studies could inform therapeutic regimens in the future. In a contrasting, but a conceptually consistent scenario, high levels of BMP antagonists such as gremlin have been reported in cancer[48], which might explain the loss of BMP responsiveness and tumor suppressive function sometimes seen in OVCA. Hence, antagonists of TGF-β/BMPs and inhibitory SMADs (SMAD6) in should be evaluated in depth in OVCA.

The importance of our finding that Sox2 is a centrally regulated target should be emphasized given that Sox2, a pioneer transcription factor[49] is overexpressed and can predict survival and prognosis in multiple cancers including ovarian[25, 26, 36, 50, 51]. Multiple epigenetic mechanisms have been shown to regulate Sox2 expression[52], and notably, these are exploited by the SMADs, as demonstrated here . Both increased promoter methylation at the dense CpG islands and H3K27me3/H3K4me3 enrichment occurred on the *SOX2* gene in a SMAD signaling and TGF-β ligand dependent manner. Interestingly, H3K27me3 and H3K4me3 enrichment occurred in the same regions where SMAD1 and SMAD3 *SOX2* gene occupancy were detected, in response to BMP9 and TGF-β1 respectively. The contribution of DNA methylation in response to BMP9 is significant and in conjunction with H3K27me3, this likely explains the high level of Sox2 repression observed in response to BMP9.

Despite the known significance of Sox2, the precise function in OVCAs has been challenging to pin down, likely in part due to the wide range in expression levels observed in cell lines (Fig. 2c). We believe that a major reason for this is the highly sensitive and context-dependent regulation of Sox2, as demonstrated here. Specifically, we demonstrate that Sox2 expression is consistently increased in response to culture under anchorage-independence and highly responsive to regulation by TGF-β family members. These findings suggest that both intrinsic cellular states and the growth factor tumor microenvironment strongly influence Sox2 regulation and may ultimately impact the effect of Sox2 perturbation as well.

Our findings delineate the specific molecular machinery utilized by TGF-β superfamily members TGF-β and BMP that determine their divergent functions on tumor cell anoikis via the central player Sox2. This study provides new information on the impact of changing the balance in growth factors in the ovarian cancer ascites environment, which will inform targeting of these pathways for therapeutic approaches to suppress ovarian cancer metastasis and tumor progression and improve patient outcomes.

### Experimental Procedures

*Cell Lines, Antibodies and Reagents (*summarized below in Table 1.0.)

**Table 1.0.** Key Resources Table

| REAGENT | SOURCE | IDENTIFIER |
| --- | --- | --- |
| **Cell lines** |  |  |
| PA1 | ATCC | CRL-1572 |
| SKOV3 | ATCC | HTB-77 |
| HEY | ATCC | N/A |
| HEK293 | ATCC | CRL-1573 |
| OVCAR3 | NIH | NCI60 (0507709) |
| OVCAR4 | NIH | NCI60 |
| OVCAR5 | NIH | NCI60 |
| M41 | Susan Murphy | N/A |
| IGROV | NIH |  |
| A2780 | Susan Murphy | N/A |
| OVCA420 | Susan Murphy | N/A |
| OVCA433 | Susan Murphy | N/A |
| OVCA429 | Susan Murphy | N/A |
| SKOV3-luc-GFP | Lab Stock | N/A |
| PA1-Luc-GFP | Lab stock | N/A |
| p76 | Amir Jazaeri | N/A |
| p211 | Amir Jazaeri | N/A |
| BON-1 | Renata Jaskula-Sztul | N/A |
| H727 | ATCC | NCI-H727 |
| A549 | ATCC | CRM-CCL-185 |
| EOC15 | Penn State and Lab stock | N/A |
| AF68 | UAB | N/A |
| **Commercial Kits** |  |  |
| Mini/Midi prep Kit | Zymo Research | D4036 |
| LIVE/DEAD™<br>Viability/Cytotoxic kit | Fisher Scientific | L3224 |
| MethylAmp DNA<br>modification kit | Epigentek | P1001 |
| DNeasy Blood & Tissue<br>kit | Qiagen | 69504 |
| **Antibodies** |  |  |
| phospho-SMAD1/5 | Cell Signaling Technology<br>(CST) | AB_491015 |
| phospho-SMAD2/3 | CST | AB_2631089 |
| SMAD1 | CST | AB_10858882 |
| SMAD2/3 | CST | AB_10693547 |
| H3K27me3 | CST | AB_2616029 |
| H3K4me3 | CST | AB_2616028 |
| β-actin | CST | AB_2242334 |
| Sox2 | CST | AB_2195767 |
| Normal Rabbit IgG | CST | AB_1031062 |
| HA-tag | CST | AB_10693385 |
| **Other reagents and recombinant constructs** |  |  |
| BMP2,4,9,10,TGFβ,<br>Activin | R&D | 355-BM, 314-BP, 3209-BP, 2926-BP, 240-B |
| Luciferin | Goldbio | LUCK |
| Dorsomorphin | Sigma-Aldrich | p5499 |
| SB431542 hydrate | Sigma-Aldrich | S4317 |
| 5-Azacytidine | Sigma-Aldrich | A2385 |
| LDN193189 | Sigma-Aldrich | SML0559 |
| ML347 | Tocris Bioscience | 4945 |
| Cycloheximide | VWR | 97064-724 |
| Actinomycin D | Fisher Scientific | AAJ60148LB |
| MG132 | Fisher Scientific | 17485 |
| shSMAD1 lentivirus | Dharmacon | V3SH11240 |
| NTC lentivirus | Dharmacon | VSC11709 |
| siSMAD4 | OriGene | SR320899 |
| siScramble | OriGene | SR30004 |
| siSMAD3 | Ambion | 4392420 |
| siSox2 | Ambion | AM16708 |
| siNTC | Ambion | 4390843 |
| Plasmid: pCMV3 | Sino biological | CV011 |
| EF1A-SOX2 lentivirus | Cellomics Tech | PLV-10013 |
| LV-CMV-SOX2 lentivirus | Cellomics Tech | PLV-10008 |
| shSox2 | Sigma | TRCN355694, TRCN3253 |
| Plasmid: pGL3-Sox2 promoter | Addgene | 101761 |
| LucGFP | Addgene | 39196 |
| Adenovirus: pc3-ALK3QD-HA | Miyazono[61] | N/A |
| Adenovirus: pc3-ALK2QD- HA | Miyazono[61] | N/A |
| Plasmid for renilla | Lab Stock | N/A |
| Mach4 Universal Detection Kit | BioCare | # M4U534 |
| Background Punisher | BioCare | #BP974 |
| Da Vinci Green Diluent | BioCare | #PD900 |
| 3,3'-diaminobenzidine (DAB) | BioCare | #BDB2004 |
| TUNEL kit | ABP Biosciences | A049 |
| PureLink PCR Purification Kit | Invitrogen | #K310002 |
| Alanine Transaminase (ALT) assay kit | Cayman Chemical | 700260 |
| Human BMP9 DuoSet ELISA | R&D | DY3209 |
| Matrigel | Fischer Scientific | 47743-720 |
| SRB salt | Aesar | A14769-14 |
| Lipofectamine RNAimax | ThermoFisher Scientific | 13778075 |
| Lipofectamine LTX | ThermoFisher Scientific | 15338100 |
| Opti-MEM medium | ThermoFisher Scientific | 31985070 |
| Dual Luciferase Assay System | Promega | E1910 |
| **Mouse strains used** |  |  |
| NOD/SCID | Jackson Labs | 001303 |
| Fox Chase SCID | Charles River | CB17/lcr |
| **Softwares used** |  |  |
| ImageJ | NIH | https://imagej.net/ImageJ |
| Prism9 | GraphPad | https://www.graphpad.com/scientific-software/prism/ |
| Adobe Illustrator | Adobe | https://www.adobe.com/products/illustrator.html |

Authentication was carried out at UAB’s Heflin Center for genomics by STR profiling. Cell lines were culture in RPMI-1640 (ATCC® 30-2001^TM^) containing L-glutamine, 10% fetal bovine serum (FBS), and 100U of penicillin-streptomycin except OVCAR3, which were cultured with 20% FBS. Patient ascites-derived EOC15 and AF68 cells were culture in 1:1 MCDB 105 and MCDB 131 with 15% FBS. HEK293 cells were maintained in complete DMEM supplemented with L-glutamine, 10% FBS, and penicillin-streptomycin. All cell lines were maintained at 37°C in a humidified incubator at 5% CO_2_, routinely checked for mycoplasma (MycoAlert PLUS mycoplasma detection kit, Lonza, Basel, Switzerland), and experiments conducted within 3-6 passages of testing depending on the cell line. Luc-GFP cell lines were generated using pHIV-Luc-ZsGreen construct. PA1 and SKOV3 cells were transduced followed by cell sorting at the UAB Flow Cytometry Core to generate stable PA1-Luc-GFP and SKOV3-Luc-GFP cells.

#### RNA interference and over-expression

Sequences for all constructs and primers are in Tables 1.0-4.0. Lentiviral particles were generated as previously described[27]. For stable SMAD1 knockdown, cells were infected with a pool of three individual SMAD1 shRNA lentivirus or non-targeting control (NTC) constructs in complete RPMI media. The media was changed after 24 hr to fresh RPMI supplemented with 10% FBS and 1×PS and left for an additional 48 hr. siRNA-mediated knock-down of SMAD3 and Sox2 was achieved using a pool of two independent siRNA duplexes to SMAD3 or Sox2, respectively and a scrambled siRNA duplex used as a negative control. Transfection was performed using Lipofectamine RNAimax reagent. Briefly, 1 × 10^5^ cells were cultured in 6 well plates in full serum medium for 24 hours. Medium was replaced with 1 ml Opti-MEM, containing 10 nM siRNA duplexes and 7.5 µl Lipofectamine RNAimax. After 15-24 hours, 1 ml 10% serum medium was added to the cells and incubated for 72 hours. The knockdown was confirmed by qRT-PCR (sequence in Table 2.0) and / or western blotting. For adenovirus infection, cells were infected with 100 MOI of adenovirus construct expressing ALK2 (Q-D)-HA, ALK3 (Q-D)-HA, generously provided by Gerard C. Blobe and Miyazono K. Transient DNA transfections were carried out in PA1 cells using Lipofectamine LTX dissolved in Opti-MEM medium. For Sox2 stable overexpression cell lines, indicated cells were infected with EF1A-Sox2 and LV-CMV-Sox2 lentivirus and their respective controls (Table 1.0) independently in complete RPMI media with polybrene for 24 hr per instructions. The media was then changed to fresh growth media and incubated for 48 hr, followed by puromycin selection.

**Table 2.0.**
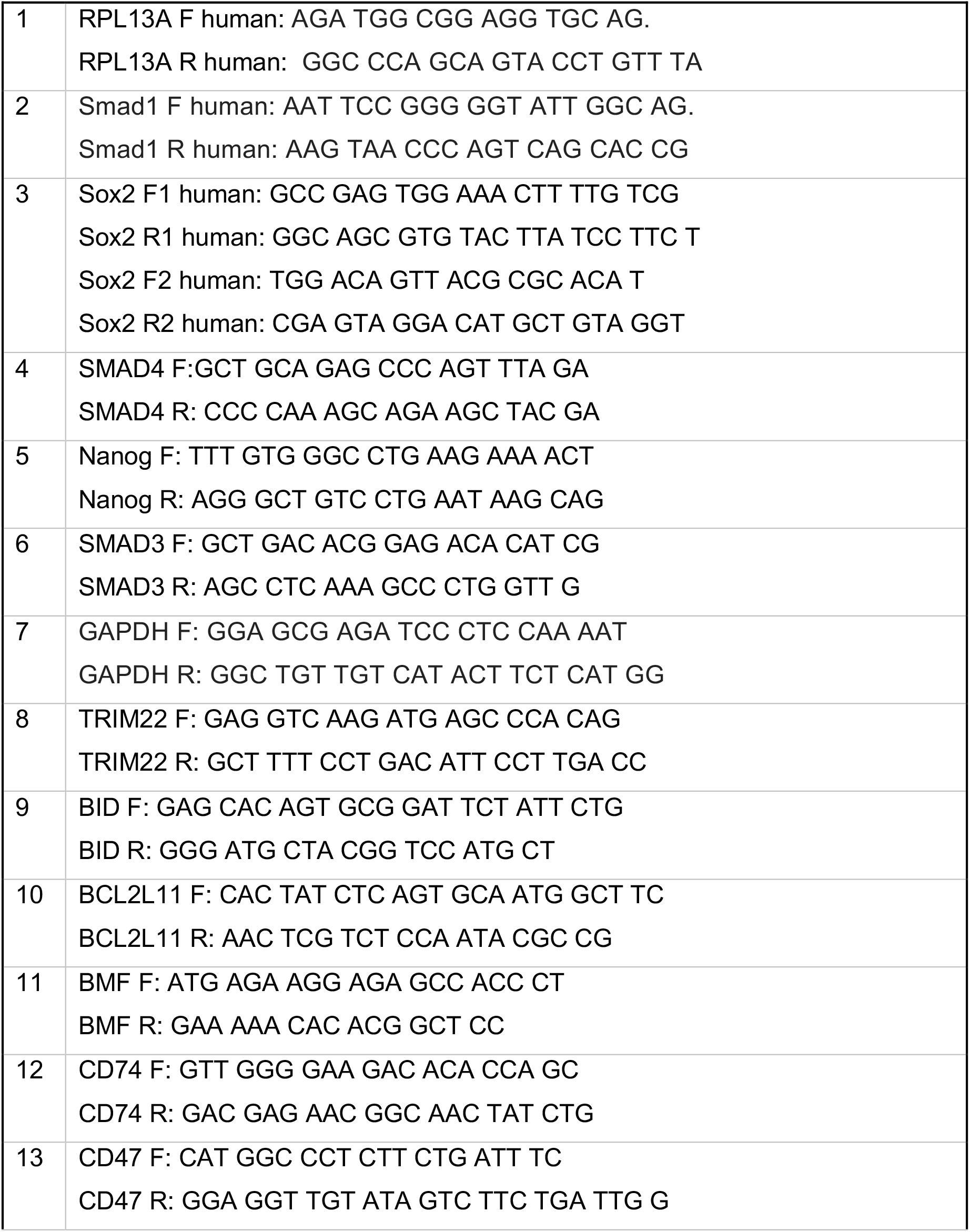
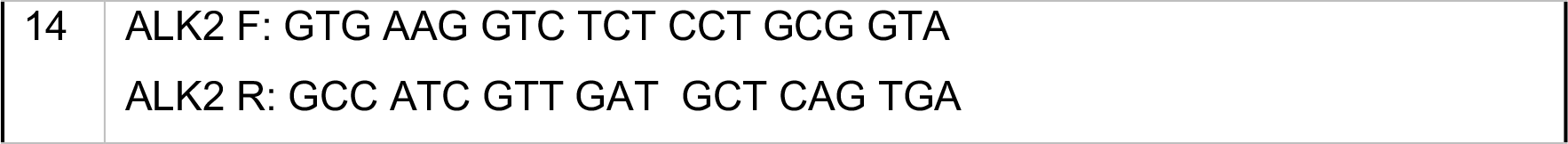
qRT-PCR Primers (listed 5’ to 3’)

#### Anchorage-independence anoikis assays

For Live/Dead analysis under anchorage independence, 1,000 cells were seeded in 96-well hydrogel-coated ultra-low attachment (ULA) plates (Corning #4515) for times indicated. Cells were stained with 2 µM Calcein-AM and 4 µM Ethidium-homodimer for 30 min before imaging. z-stacked images were obtained using the Zeiss LSM700 Confocal microscope (Microscopy and Flow Cytometry Core, University of South Carolina) and NIKON A1 Confocal microscope (UAB High Resolution Imaging Facility). Fluorescent quantification was performed using ImageJ Fiji software to calculate the Corrected Total Cell Fluorescent (CTCF) = Integrated Density – (Area of selected cell x Mean fluorescence of background readings) per spheroid.

Lysate and RNA preparation: 100,000-300,000 cells were seeded in a poly-HEMA coated 6-well plate for indicated times in full serum unless indicated otherwise. Cells were collected by centrifugation and lysed with trizol for RNA extraction or 2x lysis buffer for protein lysates.

#### 3D spheroid invasion

Spheroid invasion through Matrigel was performed as previously described[53]. Specifically, Matrigel was mixed with BMP2 and BMP9 to ensure a final concentration of 10nM and allowed to solidify for 1hr followed by addition of growth media with BMP as the top layer.. Invasion was monitored for up to 120 hr. Quantification of the amount of spread/invasion was done using ImageJ software.

#### SRB Growth Assay

Growth of cells was monitored by seeding 10,000 cells per well in a 96 well plate, followed by treatment with 10 nM BMP2/9 for indicated times. At endpoint, medium was removed from wells and SRB assay conducted as described previously [54] and absorbance measured at 570 nm with a plate reader.

#### Animal Studies

All mouse studies were performed in accordance with the Institutional Animal Care and Use Committee at the University of Alabama Birmingham. Female SCID mice (Table 1.0) at 5–7 weeks of age were housed under pathogen-free conditions at the Animal Research Facility at UAB. 1.5×10^6^ GFP-luciferase SK-OV3 cells or 3×10^6^ GFP-luciferase expressing PA1 cells were intraperitoneally injected. Mice were monitored daily with girth and weight measurement taken twice a week. Tumor progression was tracked weekly using the IVIS Lumina III In vivo Imaging System (Caliper Life Sciences, MA) at UAB’s Small Animal Imaging Facility. rhBMP9 (Table 1.0) was administered at the time of tumor cell injection followed by daily intraperitoneal (i.p.) injections of 1 mg/mL in 4mM HCl + PBS + DI water (Vehicle), to achieve a final 5 mg/kg dose. For metastasis and tumor growth analysis, mice were euthanized between 21-50 days depending on the cell line. At necropsy, ascites, if present, were collected and volumes measured, tumor weights in the omentum and other organs were recorded and collected when possible. For survival studies, mice that reached end-point criteria, including continued weight loss, respiratory trouble and permanent recumbency were euthanized. For microscopic analysis of tissues, formalin-fixed tissues were processed, paraffin-embedded, and sectioned at 5 μm thickness and H&E stained at UAB’s histology core

#### Immunohistochemistry (IHC) and TUNEL assay

IHC was performed using the BioCare Mach4 Universal Detection Kit. Specifically, anti-Sox2 was diluted in Da Vinci Green Diluent and incubated overnight at 4°C in a humidified chamber. HRP was detected with 3,3′-diaminobenzidine (DAB) substrate for 4 minutes. TUNEL staining was performed according to the manufacturer’s instruction. Slides were examined and images captured with EVOS M7000 microscope. Cell profiler and Image J Fiji software were used for image quantification.

#### Microarray and RNA sequencing

Total cellular RNA was extracted using Trizol reagent according to the manufacturer’s protocol. RNA quality was determined using an Agilent 2100 Bioanalyzer and an RNA 6000 Nano kit (Agilent, Cat. No. 5067-1511) with RNA integrity numbers (RIN) ranging from 9.8 to 10. Microarray analysis were performed on the GeneChip™ Human Gene 2.0 ST ArrayS (Thermo Fisher Scientific, Cat. No. 902112) by the functional genomic core at University of South Carolina. Data were imported into the Affymetrix GeneChip Expression Console 1.4.1.46 and processed at the gene-level using the Robust Multichip Analysis (RMA) algorithm to generate CHP files. Experimental-group specific transcriptional responses were determined using unpaired one-way between-subject analysis of variance (ANOVA). Differentially expressed genes with p-values smaller than 0.05 and fold change higher than 2.0 and lower than -2.0 were used for further bioinformatics analysis.

For RNA sequencing: library preparation was performed on purified, extracted RNA using a KAPA mRNA HyperPrep Kit (Kapa, Biosystems, Wilmington, MA) according to the manufacturer’s protocol. High throughput sequencing with 75-bp single-end reads was performed on an Illumina NextSeq 550 using an Illumina NextSeq 500/550 High Output Kit. Reads were aligned to the human transcriptome GENCODE v35 (GRCh38.p13) using STAR and counted using Salmon[55, 56]. Normalization and differential expression analysis were performed using the R package DESeq2[57]. Genes where there were fewer than three samples with normalized counts less than or equal to five were filtered out of the final data set. Benjamini-Hochberg-adjusted p-value of p < 0.05 and log2 fold change of 1 were the thresholds used to identify differentially expressed genes between treatment conditions.

#### Primary Epithelial ovarian cancer cells (EOC’s) and patient ascitic fluid ELISAs

Cells from patient ascites with an initial diagnosis of high-grade serous adenocarcinoma were collected after informed consent at the Pennsylvania State University College of Medicine (Hershey, PA) or the University of Alabama Birmingham, with approval for the study granted from the Penn State College of Medicine and UAB Institutional Review Boards (IRB). Epithelial cancer cells were isolated from ascites, as previously described[58] and used to derive EOC15 and AF68 cells respectively. AF68 was subsequently determined to favor an upper GI primary tumor with a less likely gynecological origin. For the ELISA study, ascites from patients with a confirmed diagnosis of primary OVCA were analyzed. Ascitic fluid was collected and banked after informed consent at Duke University Medical Center, with approval for the study from Duke University’s IRB. Single plex ELISAs were carried out for TGF-β1 and TGF-β2 using Aushon Biosystems Custom Circa Chemiluminescent Array kit while BMP9 was detected as described previously[59].

#### Luciferase Assay

HEK293 cells were transfected with the pGL3-Sox2 promoter-luciferase reporter plasmid construct and SV40-renilla for 24h. Treatment with BMP2 or BMP9 or TGF-β was carried out for 24 hr in serum-free media at either 10nM or 400pM respectively. According to the manufacturer’s instruction, cells were collected and lysed in 1 × passive lysis buffer. To measure luciferase activity, 20 μl of lysate was added to 25 μl of dual Luciferase Assay Reagent, and luminescence was quantitated using a luminometer (Biotek).

#### Chromatin Immunoprecipitation (ChIP) Assay

ChIP was carried out using a modified version of a previously described protocol[60]: Briefly, cells were grown to 80% confluency in 150cm^2^ culture dishes. Chromatin was sonicated using QSonica sonicator (model CL-188) for four cycles (30% amplitude for 15secs ON and 30secs OFF) to obtain DNA fragments with a length from 150 to 300 bp. 1/10th of the supernatant was stored as input control. ChIP was performed using Protein A magnetic beads (Dynabeads, Invitrogen #10001D) to couple 3.5 μg ChIP-grade antibodies for SMAD1, SMAD3, H3K27me3, H3K4me3, or rabbit IgG antibody overnight at 4°C. DNA was purified using the PureLink Quick PCR Purification kit (Thermo Fisher Cat #: K310001) and enrichment of DNA fragments analyzed via relative quantitative RT-PCR (qPCR) using ChIP primers to specific locations (Table 3.0). Negative and positive control regions were included in all analysis.

**Table 3.0.**
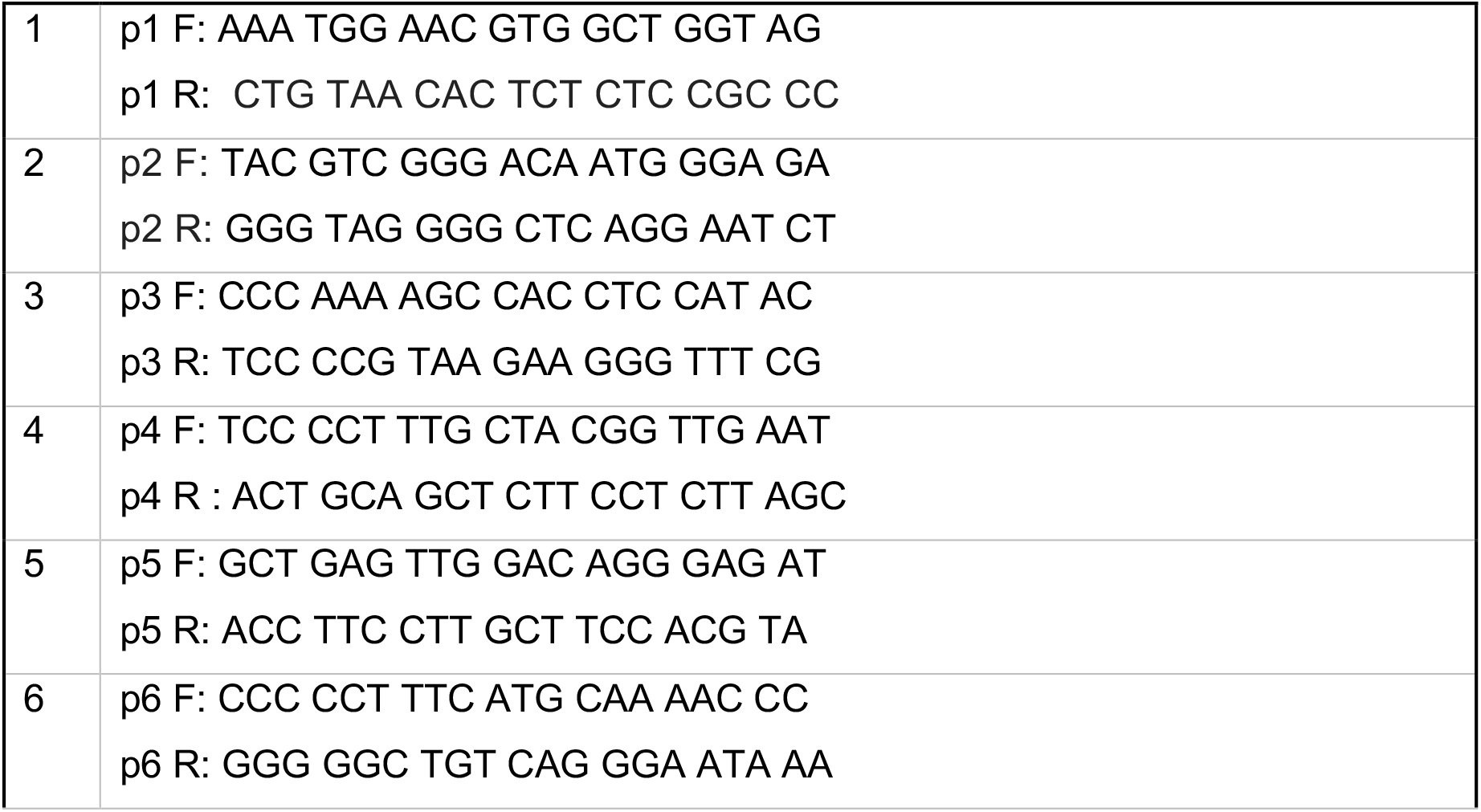
ChIP Primers

#### *Methylation-Specific* quantitative RT *PCR (MS-qPCR)*

Genomic DNA was extracted, and bisulfite conversion was performed on 500ng of gDNA using the MethylAmp DNA modification kit (Table 2.0) according to manufacturer’s instructions. Relative quantitative RT-PCR (qPCR) was performed with methylation-specific and unmethylation-specific primers (Table 4.0)

**Table 4.0.**
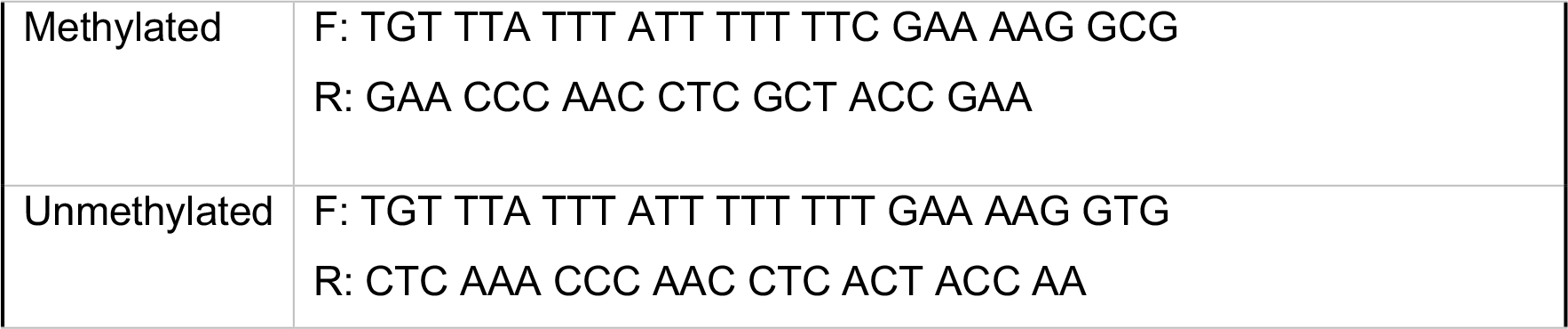
MS-PCR Sox2 Promoter primers

#### Overall Statistical analysis

Xenograft data were analyzed using parametric statistics as described in the legends. Survival curves were analyzed with log-rank statistics. *In vitro* experiments were analyzed using parametric statistics (ANOVA global test with Dunnett’s/Sidak multiple comparison test as post-hoc tests as applicable and described in legend) and presented as the mean ± SEM. All presented western blots are representative of minimum of 2-4 independent biological trials. All real time PCR’s are relative quantitative RT-PCR’s (hereby referred to as qRT-PCR) and are a combined quantitation of a minimum of 3 independent biological trials assayed in triplicate, with biological replicates represented as scatter dots in the graphs. In all cases, statistical significance was set at a threshold of p<0.05. All statistical analyses were conducted with GraphPad Prism Software.

## Acknowledgements

We would like to thank Pratik Patel, Dr. Archana Varadaraj, Dr. Lauren Vaughn, Dr. Michael Gatza and Dr. Eduardo Listik for helpful discussions and technical assistance, and Dr. Susan Murphy and Zhiqing Huang for discussions and generous gift of cell lines. Graphical abstract Images made with Biorender.com. Funding for this work was provided by NIHR01CA234969 to Mythreye Karthikeyan (KM) and Nadine Hempel (NH), American Cancer Society IRG pilot grant (University of South Carolina) and Ovarian Cancer Research Alliance’s Liz Tilberis program to Mythreye Karthikeyan (KM).

## Contributions

Z.S. contributed to study design, executed and analyzed most of the experiments and wrote the paper. M.M and K.O.C conducted a subset of experiments. S.M, assisted with cell culture studies, R.J-S provided neuroendocrine cell lines. D.A conducted and analyzed microarray studies. A.S and R.M conducted and analyzed RNA-seq studies. M.D.S and A.B.N conducted and analyzed patient ascites ELISA studies. R.P, R.A, A.B and R.W provided patient ascites. N.Y.L. assisted in data interpretation. N.H. contributed to design and data interpretation and editing of the manuscript. K.M. conceived, conceptualized, and supervised the study, designed experiments, analyzed data, and wrote the manuscript.

## Accession number

The RNA-seq data have been deposited in the NCBI-Gene Expression Omnibus (GEO) database under the accession ID GSE185932. The Microarray CEL files are accessible under Accession ID GSE185924. All the other data supporting the findings of this study are available within the article and its Supplementary information files.

