## Supplementary figures for "Reciprocal epigenetic Sox2 regulation by SMAD1-SMAD3 is critical for anoikis resistance and metastasis in cancer"

### Supplementary Fig.1

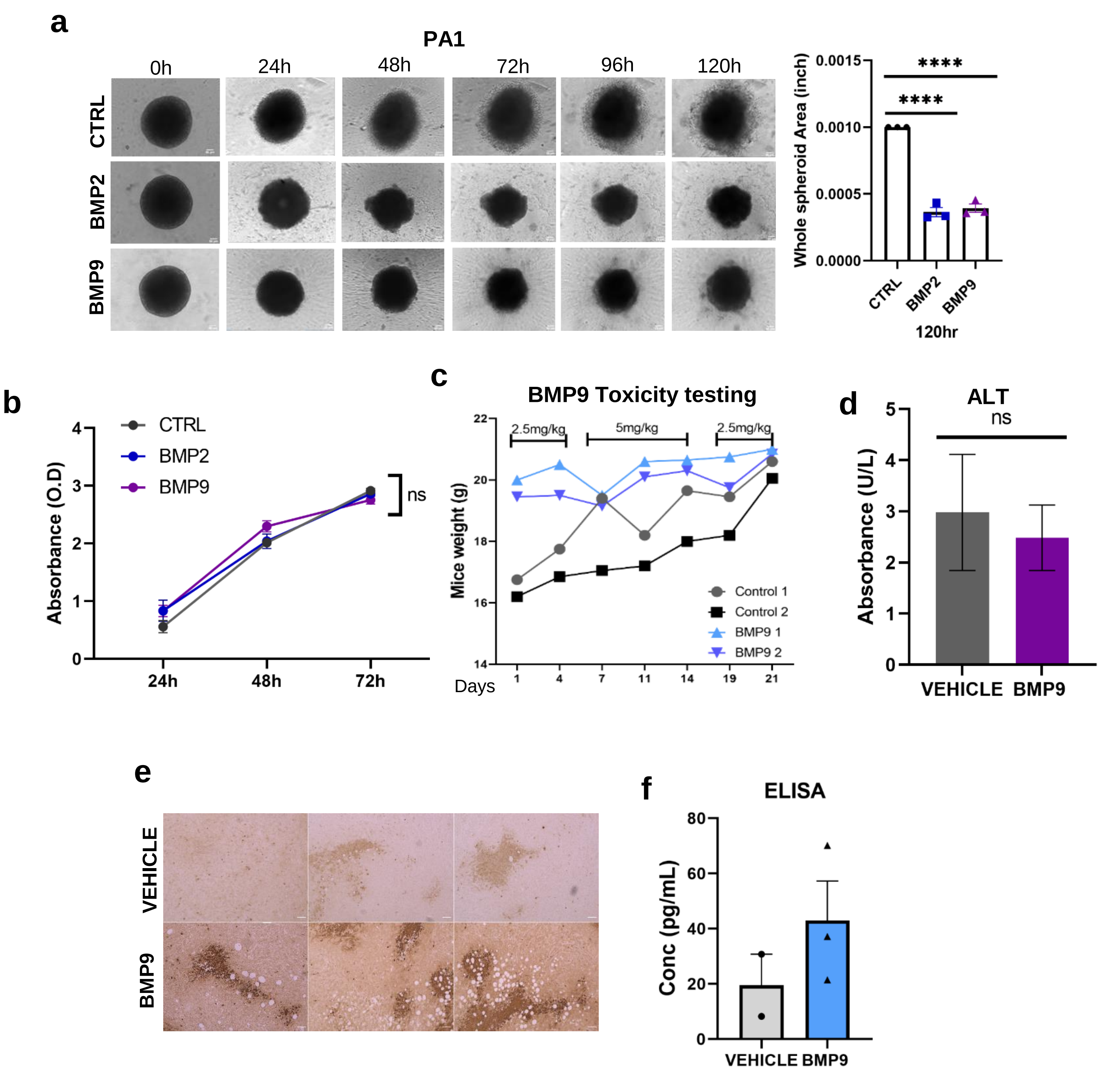


**a** 3D matrigel invasion assay of spheroids in the presence of control, BMP2 or BMP9 (10nM) (left). quantitation of spheroid invasion for the indicated time in PA1 cells (right). **b** Growth curve of PA1 cells grown under attached 2D conditions in the presence of control, BMP2 or BMP9. **c** Body weight in grams of NOD-SCID mice receiving either vehicle or BMP9 at indicated doses for a 21 day period (n=2 per group). **d** Absorbance units of Alanine Transaminase (ALT) as a measure of liver function measured from plasma (n=2 per group). **e** Representative Necrotic region in tumors from rhBMP9 vs vehicle receiving mice injected with SKOV3-luc-GFP (Scale bar= 50µM). **f** Elisa of BMP9 in plasma from mice in vehicle and BMP9 treated groups in PA1-luc-GFP mice. Data are presented as mean ± SEM, *$p$ < 0.05, **$p$ < 0.01, ***$p$ < 0.001 (**a**) ANOVA followed by Dunnett's multiple comparisons test. (**b-f**) unpaired Student's t-test

### Supplementary Fig.2

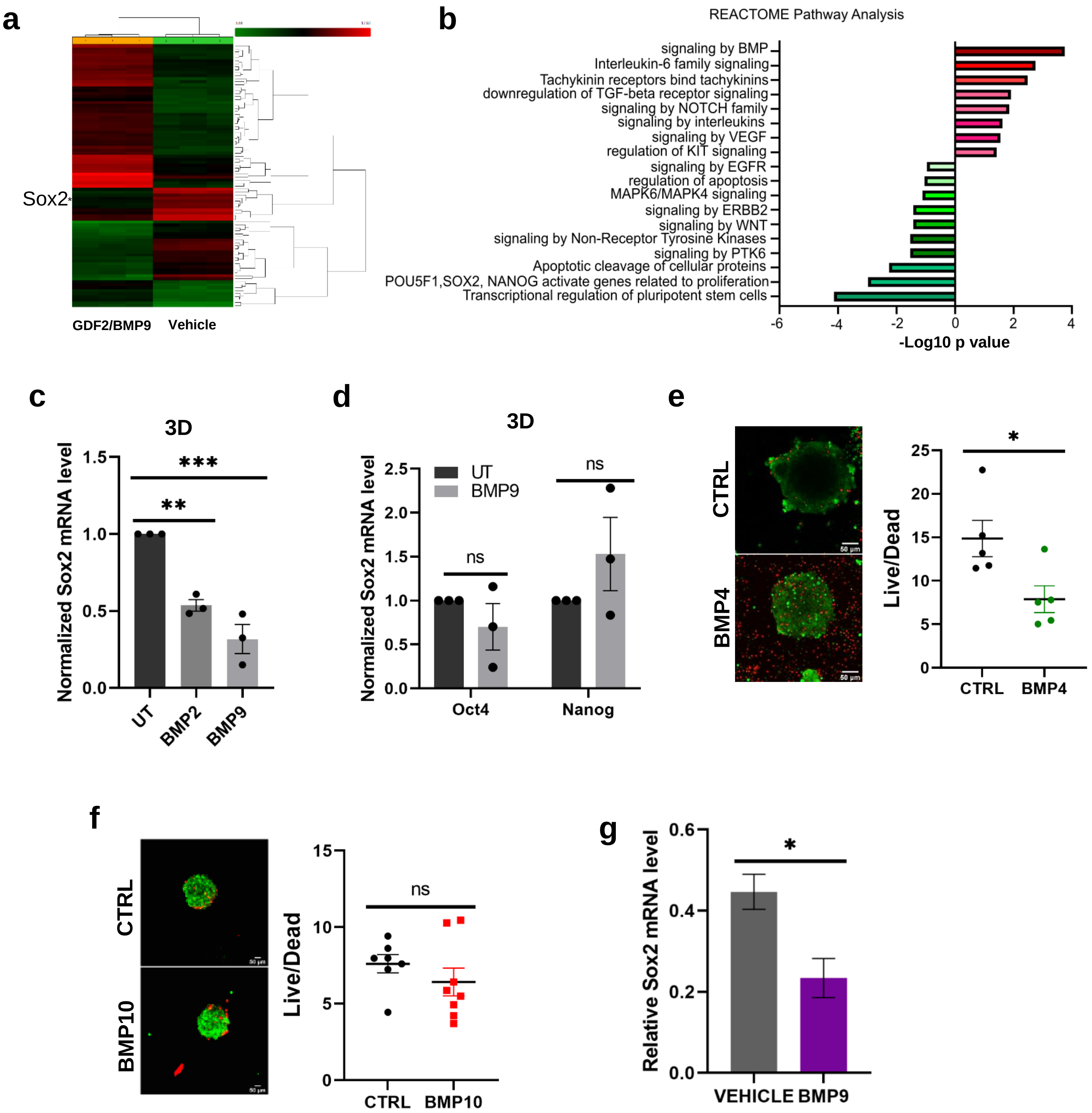


**a** Heatmap of transcription profile of 48,226 genes in PA1 cells treated with BMP9 for 24 hr under anchorage independence. **b** REACTOME pathway analysis of genes from data in (**a**). **c** Relative qRT-PCR analysis of Sox2 after BMP treatment for 24 hr under anchorage independence (3D) in PA1 cells. **d** qRT-PCR analysis of Oct4 and Nanog after BMP9 treatment under anchorage independence (3D) condition in PA1 cells. **e** Representative images of PA1 cells cultured under anchorage independence for 48 hrs, and subsequently treated with either vehicle control or with 10nM BMP4 for 24 hrs. Live/dead cell ratios were assessed by staining with Calcein-AM (green=live cells) and Ethidium homodimer dye (red= dead cells) and images taken by confocal microscopy. Scale bar = 50μm. (n=5). **f** Representative images of PA1 cells cultured under anchorage independence for 48 hrs, and subsequently treated with either vehicle control or with 10nM BMP10 for 24 hrs. (n=8). **g** qRT-PCR analysis of Sox2 in tumors from vehicle and BMP9 treated groups in SKOV3-luc-GFP mice (n=2 per condition). Data are presented as mean ± SEM, *$p < 0.05$, **$p < 0.01$, ***$p < 0.001$; (**c-d**) ANOVA followed by Dunnett's multiple comparison test; and (**e-g**) unpaired Student's t test.

### Supplementary Fig.3

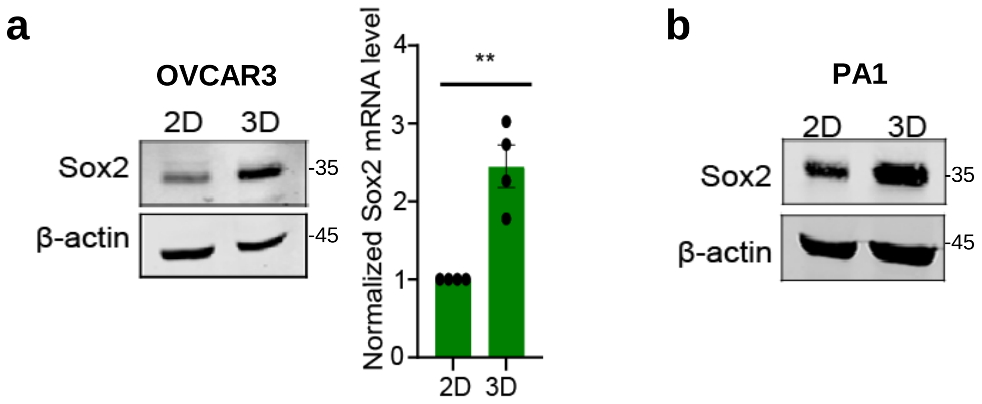


**a** Western blot (left) and relative qRT-PCR (right) of Sox2 expression under attached (2D) versus under anchorage independence (3D) conditions in OVCAR3 cells after 72 hr under 3D condition. **b** Western blot of Sox2 expression under 2D versus 3D condition in PA1 cells after 72 hr under 3D condition. Data are presented as mean ± SEM, **p* < 0.05, ***p* < 0.01, ****p* < 0.001

### Supplementary Fig.4

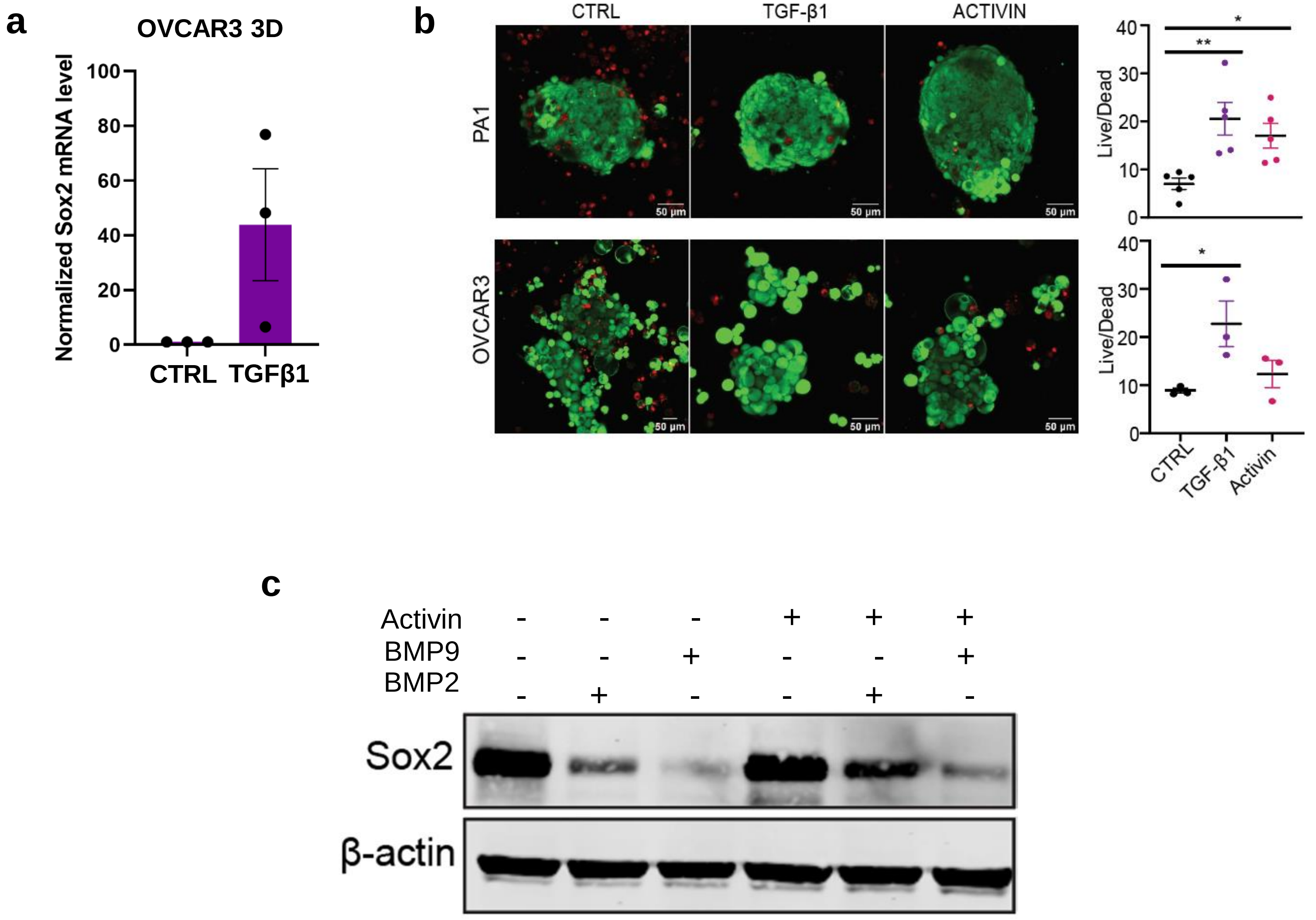


**a** Relative qRT-PCR analysis of Sox2 after TGFβ1 treatment for 24 hrs under anchorage independence (3D) condition in OVCAR3 cells. **b** Live-dead analysis of cells under anchorage independence after TGFβ1 and activin treatment for 24 hr in indicated cells (n=3 to 5). **c** Western blot of Sox2 after combined treatment of equimolar (10nM) activin and BMP2/9 for 24 hr in PA1 cells (n=2). Data are presented as mean ± SEM, *$p$ < 0.05, **$p$ < 0.01, ***$p$ < 0.001. (ANOVA followed by Sidak's multiple comparisons test).

### Supplementary Fig. 5

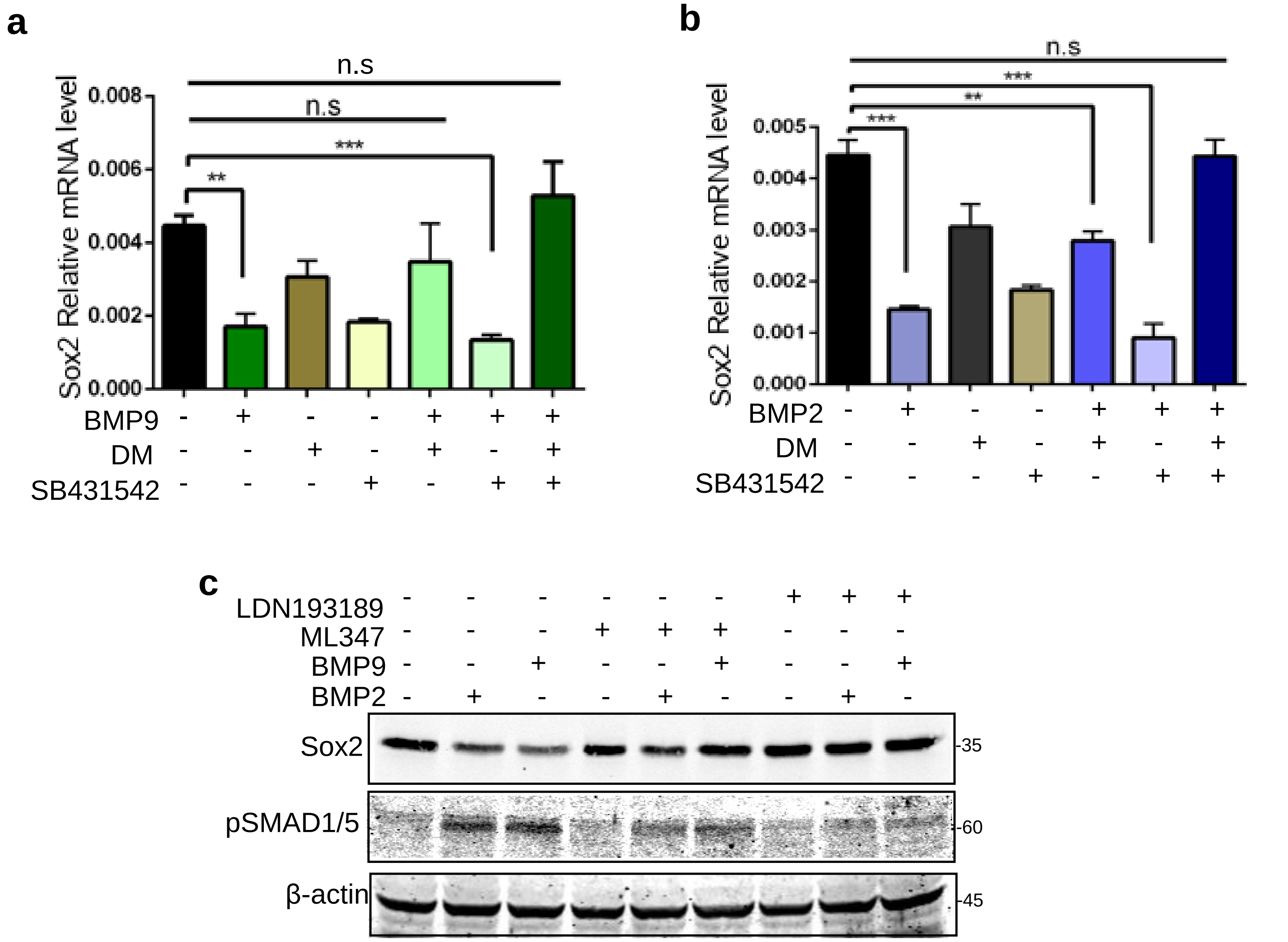


**a** qRT-PCR analysis of Sox2 expression in OVCAR3 cells pretreated with 5μM Dorsomorphin (DM) and 5μM SB431542 for 1hr, followed by treatment with BMP9 for 24 hrs. Data are normalized to DMSO vehicle controls. **b** qRT-PCR analysis of Sox2 expression in OVCAR3 cells pretreated with 5μM Dorsomorphin (DM) and 5μM SB431542 for 1hr, followed by treatment with BMP2 for 24 hrs. Data are normalized to DMSO vehicle controls. **c** Western blot of Sox2 expression in OVCAR3 cells pretreated with 3μM ALK1,2 inhibitor ML347 and 0.8μM ALK2,3 LDN193189 for 1hr, followed by treatment with BMP2/9 for 24 hrs. Data are normalized to vehicle controls presented.

#### Supplementary Fig.6

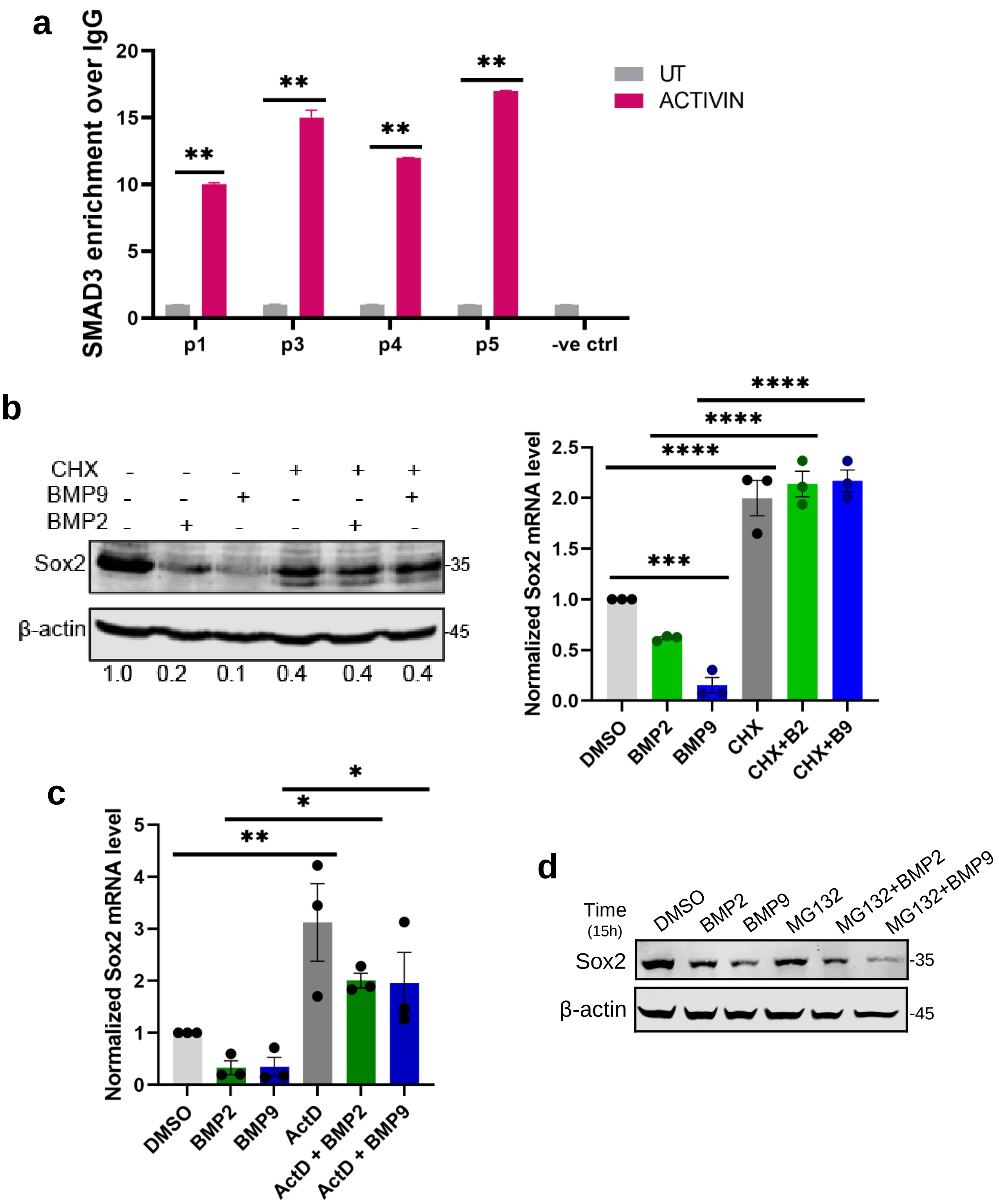


**a** Representative relative qRT-PCR of indicated regions (primer sites) after chromatin immunoprecipitation of SMAD3 to sites on Sox2 proximal chromosomal regions with or without 1hr of activin A treatment, expressed as the ratio over IgG controls normalized to untreated cells. **b** Western blot (left) and qRT-PCR (right) of Sox2 upon either 10ng/μL Cycloheximide (CHX) or BMP2 or BMP9 treatment as indicated normalized to DMSO control in PA1 cells. (ANOVA followed by Dunnett's multiple comparison test (n=3). **c** Relative qRT-PCR analysis of Sox2 in the presence or absence of 0.5ng/mL Actinomycin D (Act D) with or without BMP2 or BMP9 as indicated normalized to DMSO control for 12 hrs in PA1 cells. (ANOVA followed by Sidak's multiple comparison test). **d** Western blot of Sox2 expression in PA1 cells pretreated for 1 hr with 0.5μM MG132, followed by BMP2/9. Data are presented as mean ± SEM, **p* < 0.05, ***p* < 0.01, ****p* < 0.001.

### Supplementary Fig.7

a

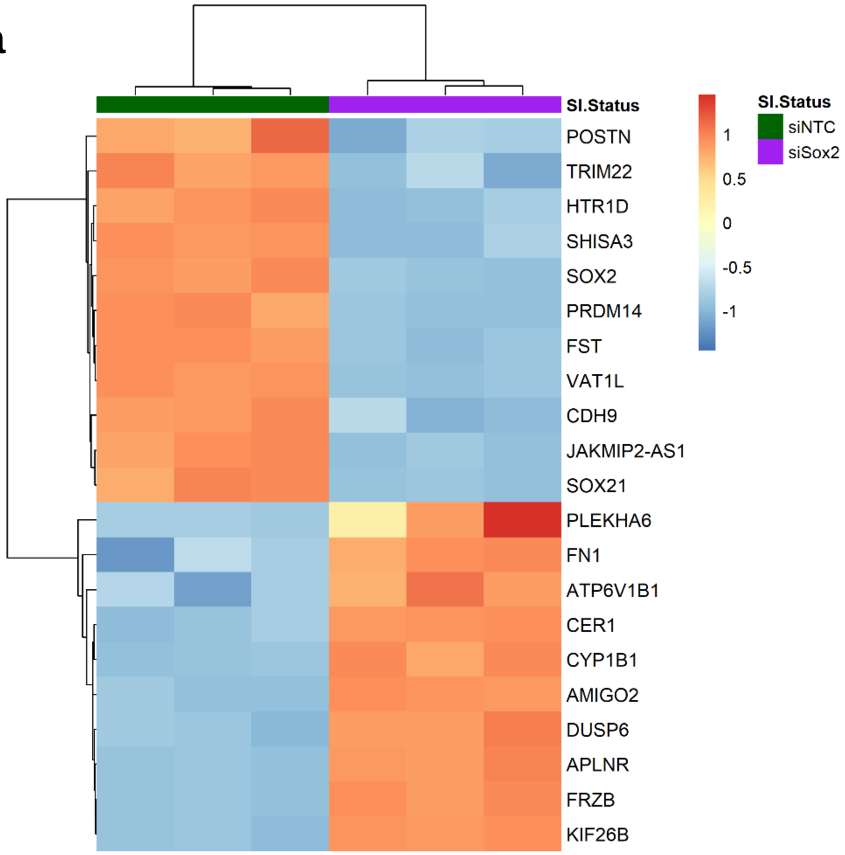


b

| GENE | LOCATION | STRAND | P-VALUE |
|---|---|---|---|
| KIF26B | 928 | - | 0.000275 |
| PRDM14 | 710 | - | 0.0001 |
| TRIM22 | 168 | - | 0.00087 |
| DUSP6 | 381 | + | 0.0001 |
| SOX21 | 404 | - | 0.0001 |
| POSTN | 218 | + | 0.0001 |
| CDH9 | 178 | + | 0.000625 |
| SHISA3 | 75 | + | 0.0013 |
| VAT1L | 261 | + | 0.000075 |
| APLNR | - | - | - |
| FN1 | 87 | - | 0.002 |
| HTR1D | 27 | + | 0.00015 |
| FST | 207 | - | 0.0006 |
| FRZD | - | - | - |
| JAKMIP2-AS1 | - | - | - |
| PLEKHA6 | 144 | + | 0.000125 |
| ATP6V1B1 | 49 | - | 0.000175 |
| AMIGO2 | 585 | + | 0.000675 |
| CYP1B1 | 358 | + | 0.0003 |
| CER1 | 116 | - | 0.0002 |

c

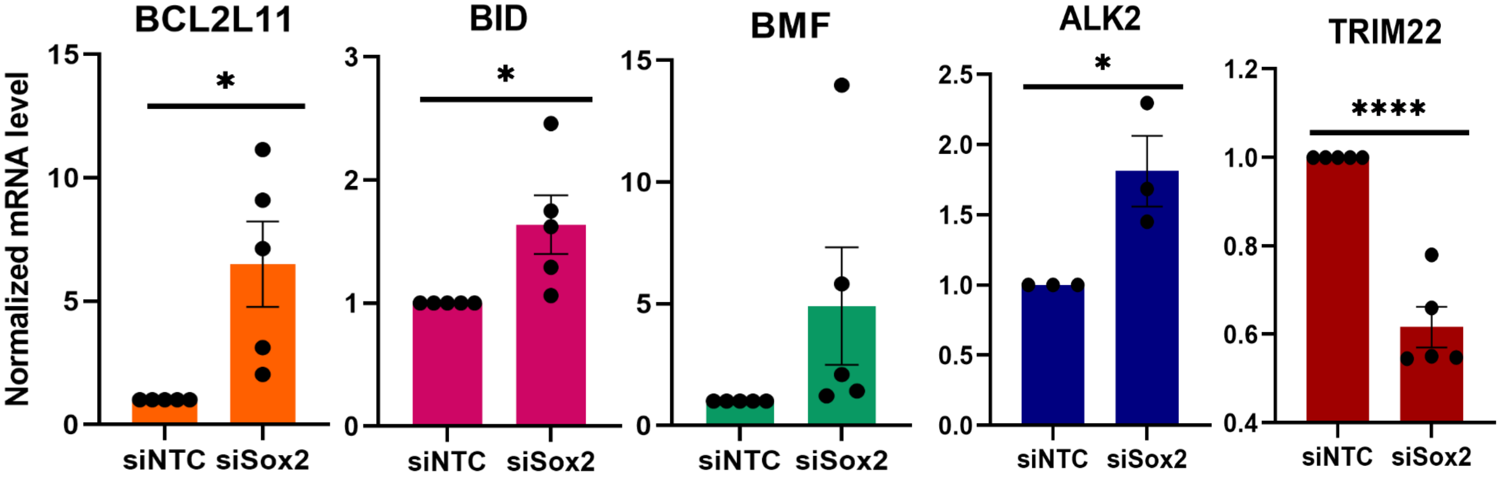


d

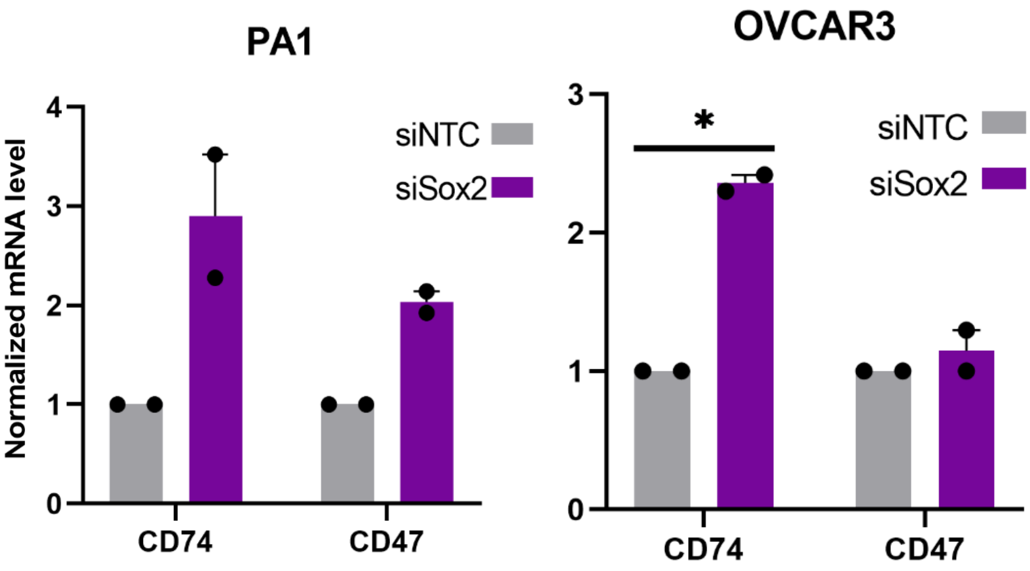

**a** Heatmap of 21 common differentially expressed genes between the microarray from GEO #GSE185924, Fig. 2a and RNA sequencing analysis GEO GSE185932, Fig. 8c. **b** LASAGNA analysis of Sox2 binding motifs for the 21 DEGs from (**a**). **c-d** qRT-PCR analysis of indicated candidate genes in siNTC and siSox2 normalized to siNTC in either PA1 cells or OVCAR3 as indicated Data are presented as mean ± SEM, *$p < 0.05$, **$p < 0.01$, ***$p < 0.001$.
